# PFKFB4 control of Akt signaling is essential for premigratory and migratory neural crest formation

**DOI:** 10.1101/168807

**Authors:** Ana Leonor Figueiredo, Frédérique Maczkowiak, Caroline Borday, Patrick Pla, Meghane Sittewelle, Caterina Pegoraro, Anne H. Monsoro-Burq

## Abstract

**Summary statement:** PFKFB4 controls neural crest final specification and migration by regulation of AKT signaling or glycolysis.

**Abstract:** Neural crest (NC) specification comprises an early phase, initiating immature NC progenitors formation at neural plate stage, and a later phase at neural fold stage, resulting into functional premigratory NC, able to delaminate and migrate. We found that the NC-GRN triggers up-regulation of *pfkfb4* (6-<u>p</u>hosphofructo-2-<u>k</u>inase/<u>f</u>ructose-2,6-<u>b</u>isphosphatase <u>4</u>) during this late specification phase. As shown in previous studies, PFKFB4 controls AKT signaling in gastrulas and glycolysis rate in adult cells. Here, we focus on PFKFB4 function in NC during and after neurulation, using time-controlled or hypomorph depletions *in vivo*. We find that PFKFB4 is essential both for specification of functional premigratory NC and for its migration. PFKFB4-depleted embryos fail activating *n*-*cadherin* and late NC specifiers, exhibit severe migration defects, resulting in craniofacial defects. AKT signaling mediates PFKFB4 function in NC late specification, while both AKT signaling and glycolysis regulate migration. These findings highlight novel and critical roles of PFKFB4 activity in later stages of NC development, wired into the NC-GRN.

## INTRODUCTION

The neural crest (NC) is a highly migratory and multipotent cell population of vertebrate embryos, forming an array of differentiated cells: peripheral neurons and glia, pigment cells, craniofacial skeleton, cardiac structures, and endocrine cells (Bronner and LeDouarin, 2012). NC development initiates during gastrulation, at the edges of the neural plate, and continues until late organogenesis. In early gastrulas, NC development is triggered by signals from the adjacent neural plate, non-neural ectoderm, and underlying mesoderm (Saint-Jeannet et al., 1997; Wilson et al., 1997; Neave et al., 1997, Chang and Hemmati-Brivanlou 1998; LaBonne and Bronner-Fraser 1998; Villanueva et al., 2002; Monsoro-Burq et al., 2003; Milet and Monsoro-Burq 2012). These signals specify the neural border (NB), a transition area located across the neural plate and the non-neural ectoderm (Meulemans & Bronner-Fraser 2004; Monsoro-Burq et al. 2005; Basch et al. 2006). In gastrulas, the NB is a mixed territory comprising prospective dorsal neural tube cells, NC and cranial placode progenitors (Steventon et al. 2009, Pegoraro & Monsoro-Burq 2012).

In late gastrulas and early neurulas (i.e. neural plate stage), NC specification is initiated within the NB, upon the coordinated action of WNT signaling and several transcription factors (NB specifiers) broadly activated in this region (PAX3/7, ZIC1/2, MSX1/2, TFAP2A, GBX2, HES4, AXUD, cMYB; Luo et al. 2003; Brewer et al. 2004; Monsoro-Burq et al. 2005; Sato et al. 2005; Khadka et al. 2006; Basch et al. 2006; Nichane et al. 2008; Li et al. 2009; Maczkowiak et al. 2010; de Crozé et al. 2011; Simoes-Costa et al. 2015). This initial phase of NC specification is marked by locally enhanced expression of *tfap2a* and *hes4/hairy2*, and low-level *snail2* and *foxd3*, two transcription factors marking the early NC progenitors, i.e. lineage-restricted cells being still functionally immature (Saint-Jeannet et al. 1989; Essex et al. 1993; Nieto et al. 1994; Mayor et al. 1995; Mancilla and Mayor, 1998; Nichane et al., 2008; de Crozé et al., 2010). During the second half of neurulation, as neural folds elevate, the immature NC is further specified into functional premigratory NC, ready to undergo epithelium-to-mesenchyme transition (EMT) and migration. The fully specified NC (mature) is characterized by late markers *sox10*, *twist* and *tfap2e*, and enhanced *ets1* and *cmyc* (Bellmeyer et al. 2003; Théveneau et al. 2007). In late neurulas, after EMT, NC cells migrate as groups or as individual cells along defined embryonic routes, still expressing *sox10* and *twist* (Theveneau & Mayor 2012). Importantly, TWIST is essential for the coordinated action of EMT regulators SNAIL1/2 (Lander et al. 2013) while enhanced survival properties favors the NC extensive migration (Vega et al., 2004). Finally, the cranial NC populating the craniofacial buds differentiates into mesenchyme, cartilage and bone (Douarin & Kalcheim 1999).

We found that *pfkfb4 (6*-*phosphofructo*-*2*-*kinase/fructose*-*2*,*6*-*bisphosphatase)* expression was upregulated during late NC specification/maturation phase (neural fold stage). In adult cells, PFKFB1-4 control the levels of fructose-2,6-bisphosphate, the strongest allosteric activator of phosphofructokinase1, which regulates the rate-limiting reaction of glycolysis (Okar et al. 2001; Pilkis et al. 1995, Fig. S1). Higher PFKFB4 levels stimulate glycolysis rate. Moreover, in embryos, we have previously shown that PFKFB4 is essential for global ectoderm regionalization. In this context, PFKFB4 acts independently of glycolysis by controlling AKT signaling (Pegoraro et al. 2015). Here, we found that the NC-GRN controls elevated *pfkfb4* expression in the late premigratory NC. Using either constitutive or time-controlled PFKFB4 depletions, and time-controlled pharmacological inhibition of PI3K-AKT signaling *in vivo*, we show that PFKFB4 and AKT signaling regulate a timely switch between lineage-restricted but immature NC and functional premigratory NC. Furthermore, after EMT, PFKFB4, through PI3K-AKT signaling and glycolysis, controls NC migration. These results suggest that NC developmental progression relies upon continuous PFKFB4 function, as a novel NC-GRN actor, whose activity is relayed by AKT signaling.

## RESULTS

### *Pfkfb4* expression labels the neural crest during second half of neurulation

As in all vertebrate embryos, in *Xenopus laevis*, NB induction starts in early gastrulas (st.10.5 Nieuwkoop-Faber, 1994), as characterized by early expression of *pax3*, *zic1/2*, and *hes4* lateral to the neural plate (Monsoro-Burq et al., 2005; Nichane et al., 2008). NC specification is initiated at the end of gastrulation (st. 11.5-12), and is marked by early NC specifiers as *snail2* and *foxd3* at neural plate stage (st. 12-14, Fig. 1A). NC specification is further established (maturation phase) during neural fold stage (st. 14-18), by activation of a later set of NC specifiers and EMT inducers (*sox10*, *twist)*, and by increased expression of early NC specifiers (Fig. 1B). Upon neural tube closure (st. 17-19), cranial NC cells undergo EMT and initiate migration to populate the branchial arches and craniofacial areas around tailbud st.24 (Fig. 1C).

**Fig. 1.**
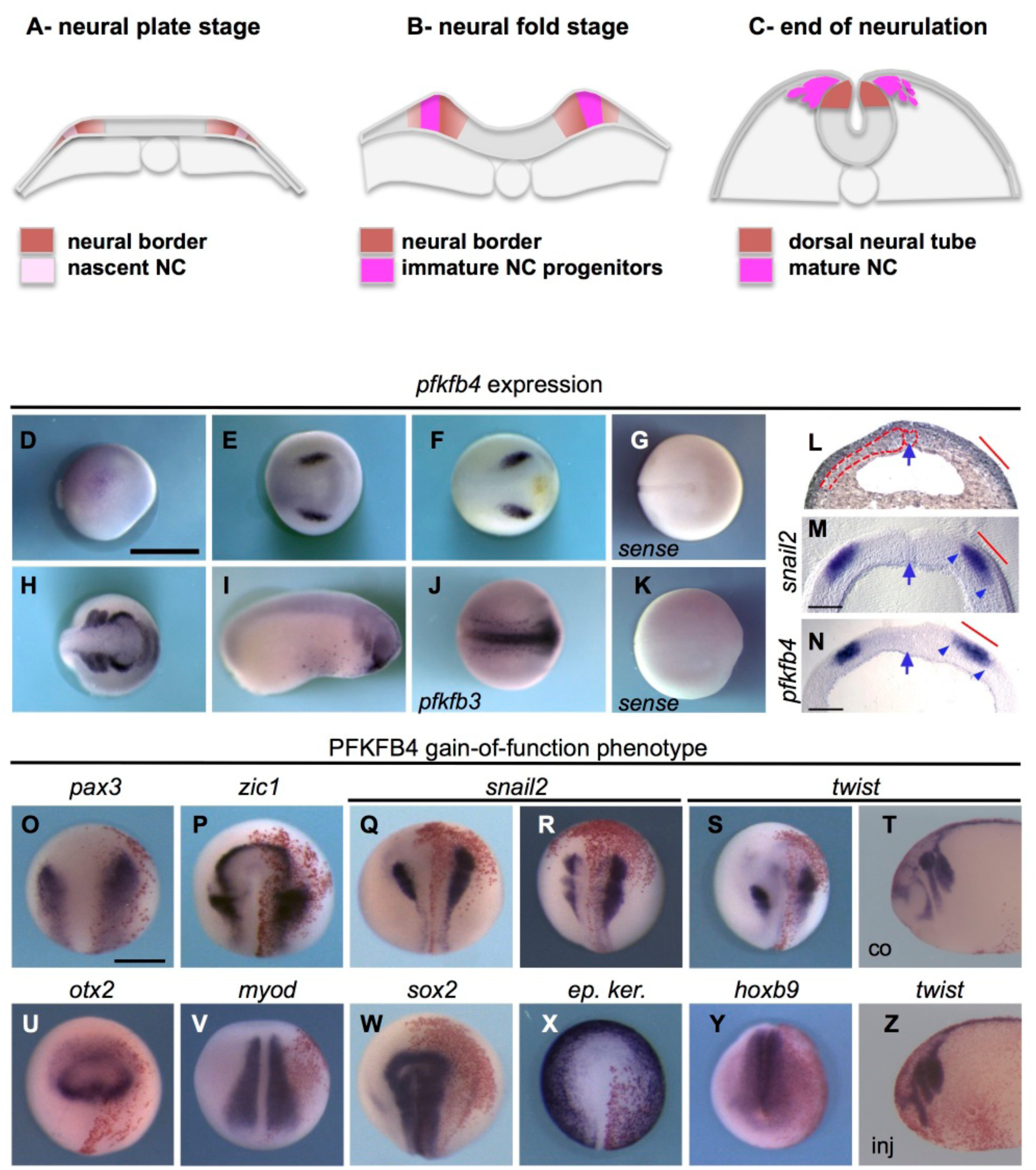
PFKFB4 promotes neural border and neural crest formation. (A) NC is induced during neural plate stage (st.12.5 to st.14). NB specifiers expression (red) is robustly established lateral to the neural plate, early NC specifiers expression (*snail2*, *foxd3)* is faintly initiated (light pink). (B-C) During neural fold elevation (st.14 to st.17), future NC cells progressively acquire their definitive specification and specific cellular properties (survival, expression of late NC specifiers, cadherin switch…) ensuring their ability to undergo EMT and migration at the end of neurulation (C, st.18 to st.19). (D-F,H,I) *Pfkfb4* is enriched in the NB/NC in neurulas and tadpoles. (J) *Pfkfb3* is expressed in the neural tube. Sense probes (*pfkfb4*, G; *pfkfb3*, K). Neurula stages: late gastrula/early neural plate stage: st.12 (D), end of neural plate stage, st.14 (E), neural fold stage; st.16 (F), neural tube closure/ end of neurulation: st.19 (H). Tailbud stage: st.22 (I). E-H, J,K: dorsal views, anterior to the right. D,I: side views. Scale bar, 1mm. (L-N) Cross-sections through the mid-neurula anterior neural plate (st.14) show *snail2* and *pfkfb4* expression in NB/NC. (L) The notochord, neural plate and paraxial mesoderm are outlined on hematoxylin-eosin stained sections. (M-N) Vibratome sections. Arrow indicates the midline. Arrowheads indicate the st.14 NB (red bar). Scale bar, 200μm. (O-S) *pfkfb4* gain-of-function expanded the st.14 NB (O,P, 59% of embryos with enlarged NB compared to the contralateral uninjected side, n=81) and premigratory NC (Q, st.14, 59% enlarged *Snail2* expression, n=147), (R,S: st.18, 55% increase, n=271). Migrating NC streams were expanded (T,Z, st. 22-24, 65% expansion, n=49). In contrast, pan- or regional neural plate markers (*sox2*, *otx2*, *hoxb9*), ectoderm (*keratin K81*, *ep.Ker*.) or paraxial mesoderm (*myod*) markers were unaffected (U-Y). O-S, U-X: dorsal views. T,Z: side views, Y: dorsal-posterior view. Scale bar=500μm. Detailed phenotype scoring in Table.S2. Box initially at rest on sled sliding across ice.

Among novel genes enriched in premigratory NC, we found *pfkfb4* clustered with *snail2* and *sox10* (Plouhinec et al. 2014). At neural plate stage, *pfkfb4* was expressed at low levels throughout the dorsal ectoderm (Fig. 1D). At early neural fold stage (st.14), later than *snail2* initiation, *pfkfb4* was strongly enriched in the NB/NC (Fig. 1E,F,L-N). At the end of neurulation (st.18), NC *pfkfb4* mRNA levels, but not *pfkfb1*-*3* levels, were elevated by 1.5-fold compared to whole embryo expression (Fig. 1J, Fig. S2). The premigratory NC expressed 6.4-fold higher *pfkfb4* levels than the adjacent anterior neural fold, which forms forebrain and placodes. Finally, *pfkfb4* was expressed during NC EMT and early migration but decreased as NC cells reached ventral craniofacial areas (Fig. 1H-I). This pattern contrasted with blastula and gastrula stages, when low *pfkfb4* levels were detected in the entire dorsal ectoderm, encompassing the neural plate, the NB, and part of the non-neural ectoderm (Pegoraro et al. 2015). Thus, *pfkfb4* was enriched in premigratory NC, during its maturation period, NC EMT and early migration (st14-22).

### PFKFB4 activation promotes neural crest formation at the neural border

Experimental increase of PFKFB4 levels *in vivo* led to a moderate but reproducible increase of the NB and early NC marker expression at the end of neural plate stage (st.14, *pax3*, *zic1*, *snail2*, Fig. 1O-Q, Table.S2). At the end of neurulation (st.18), the definitive premigratory NC area was also enlarged (Fig. 1R,S). In contrast, neural plate specification, anterior-posterior regionalization (*sox2*, *otx2*, *hoxb9*) and non-neural ectoderm differentiation (*keratin*-*K81*) were not affected in late neurulas. We observed a transient and moderate increase of the *sox2*-positive area at mid-neurula stage, which was probably related to the enlarged NB, which transiently expresses *sox2* early on (Fig. 1U,W-Y, Table.S2). Mesoderm formation seemed unaffected (*myod*, Fig. 1V). At tailbud stage (st.22-24), migrating NC streams were enlarged with similar ventral migration distance compared to the contralateral control side (Fig. 1T,Z). Altogether, *pfkfb4* expression and PFKFB4 gain-of-function phenotype suggested a positive role in NC development. We did not observe ectopic expression of NC markers, in the neural plate or the non-neural ectoderm, suggesting that PFKFB4 was not sufficient to alter specification of adjacent ectoderm cells.

### Moderate PFKFB4 depletion affects craniofacial development *in vivo*

In order to deplete PFKFB4 activity in gastrulas, we previously used a splice-blocking morpholino oligonucleotide (PFKFB4MO, Pegoraro et al. 2015), using 20ng per blastomere in 2-cell-stage embryos. *Pfkfb4* mRNA levels were decreased by 60% (hereafter called severe depletion). Phenotypes were validated using an ATG-targeting and a mismatch morpholino, dose-response, and rescue experiments (Blum et al. 2015). In severe depletion conditions, we observed a developmental blockade before neural fold stage, all dorsal ectoderm cells remaining in an immature progenitor status, while further specification was blocked. Here, in order to analyze later PFKFB4 functions during NC formation, we used two strategies to bypass this early developmental arrest. Firstly, we used low-level PFKFB4 depletion, which no longer blocks ectoderm global developmental progression and allows neurulation to proceed. Secondly, we have used photo-inducible morpholinos, to control PFKFB4 depletion timing *in vivo*, and avoid affecting early development.

Based on *pfkfb4* enhanced NC expression, we hypothesized that NC development required maintenance of high PFKFB4 levels, while other tissues may develop normally under mild PFKFB4 depletion conditions. Using decreasing PFKFB4MO doses, we created mild hypomorph conditions (about 35% decrease in *pfkfb4* mRNA level, Fig. 2A), and monitored the craniofacial morphology and pigmentation phenotype of st.45 tadpoles (Fig. 2B-E). We found that low-level depletion (injecting 5ng of MO into one dorsal-animal blastomere, at the 4-cell stage or any equivalent combination at other developmental stages), resulted in a morphologically normal gastrulation and neurulation but caused a significant reduction of head structures on the injected side, compared to the uninjected side (Fig. 2B). Pigment cells were present on both sides of the tadpoles and appeared unperturbed. Lower morpholino doses (2.5ng or 1ng in one of four blastomeres) did not significantly alter embryo morphology. Higher doses caused early death of the injected cells (this apoptosis was rescued by adding *pfkfb4* mRNA as described in Pegoraro et al., 2015). With the low-level PFKFB4 depletion, we checked that injected cells were not subjected to increased cell death compared to control cells in the vast majority of the embryos (Fig. S3). Moreover, neither ventral injections of PFKFB4MO nor dorsal injections of a non-related MO (even at high doses) caused cell death, excluding non-specific effects of morpholino injection (Fig. S3). These observations confirmed the hypothesis that NC development was specifically affected by low-level depletion of PFKFB4. Therefore, using these hypomorph PFKFB4 conditions, we could explore the roles of PFKFB4 in NC derivatives formation *in vivo*. All further experiments, except for photo-morpholinos, were thus conducted using this low MO concentration.

**Fig. 2.**
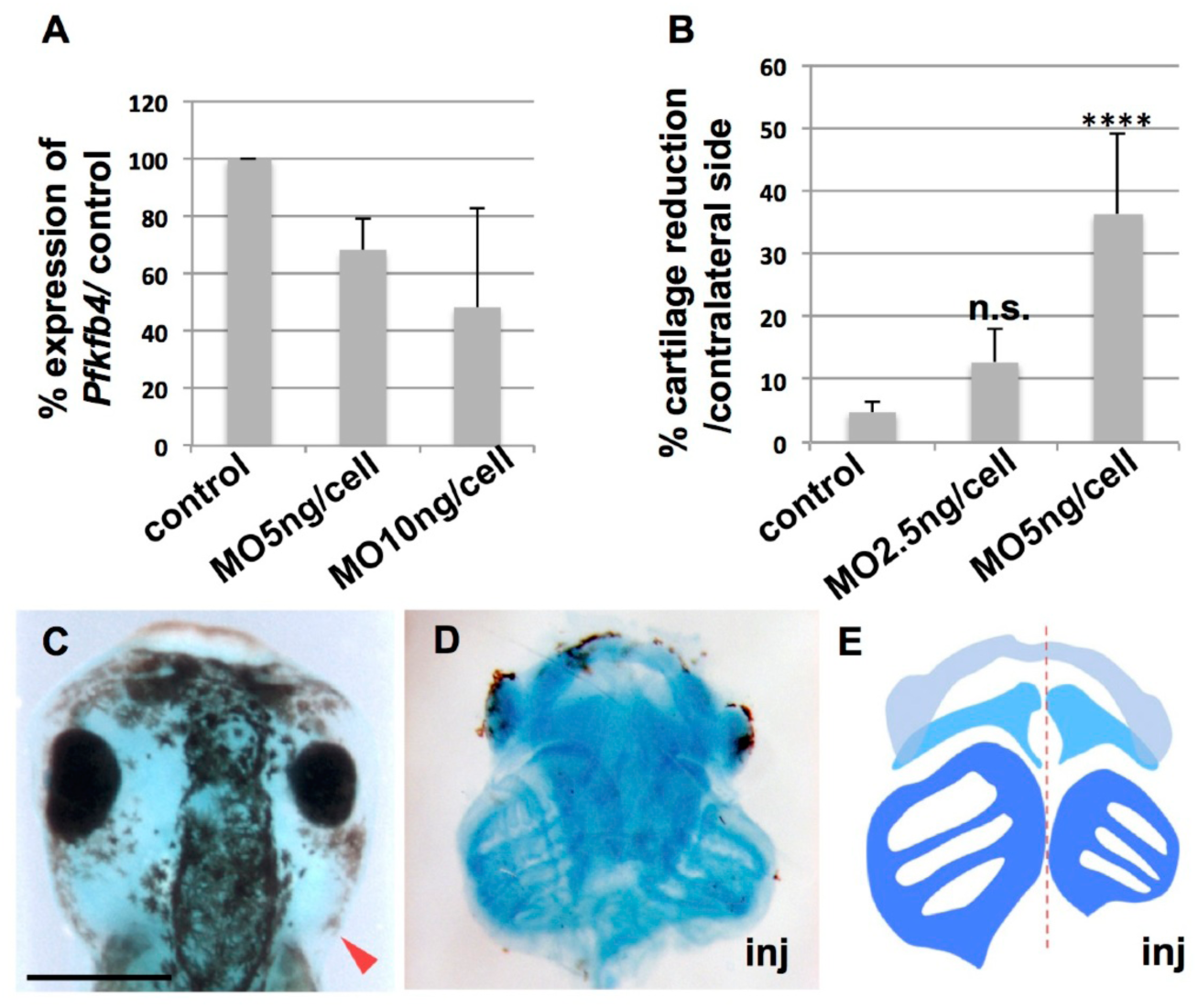
Moderate PFKFB4 depletion results in severe craniofacial defects in tadpoles. (A) *Pfkfb4* mRNA levels were quantified on neurulas dorsal tissues (RTqPCR). Injections at “high concentration” (10ng/cell in 4-cell-stage embryos) reduced *pfkfb4* levels by 52% and caused developmental arrest. “Milder” concentration (5ng/cell in 4-cell-stage embryos) decreased *pfkfb4* mRNA by 32% (low-level depletion), and allowed swimming tadpoles development. (B-E) Craniofacial morphology in tadpoles st.45-46. Low-level PFKFB4MO depletion reduced craniofacial structures on the injected side (n=17). The branchial arch cartilages area was severely diminished on the injected side (36% reduction compared to the contralateral side, n=8; p-value<0.0001 compared to side-to-side variations in a control population, n=9). Lower doses did not significantly reduce cartilage area (n= 6), or affect head morphology (n=20). (C) Head morphology, dorsal view, arrowhead: injected side. (D) Dissected jaw and branchial arches cartilages, ventral view, arrowheads: injected side. (E) Scheme of D. Scale bar,500 μm.

The NC generates most craniofacial structures including craniofacial skeleton, mesenchyme and soft tissues around the eyes. Following moderate PFKFB4 depletion, dorsal and ventral craniofacial structures were globally smaller. On the morphant side, branchial cartilages were strongly reduced while Meckel’s and ceratohyal cartilages were moderately diminished (Fig. 2C-E). Eyes were also affected, with grossly normal shape but reduced size. Since *pfkfb4* is present at low levels in the neural plate early on (Fig. 1), this could be due to a cell autonomous defect in the optic vesicles forming from the neural plate. Alternatively, CNCC defects could influence the developing eye (Bailey et al. 2006; Lwigale 2015). Hence, the NC-derived craniofacial structures were severely affected when PFKFB4 levels were downregulated moderately while most other structures were normal.

### PFKFB4 depletion affects premigratory neural crest specification and NCC migration

We then investigated which steps of NC formation were affected by PFKFB4 low-level depletion. At early neural fold stage, *snail2* expression was very low to absent on the injected side (Fig. 3A,L-first bar, Table.S2). This defect was a delay, since, by the end of neurulation, most embryos expressed *snail2* normally (Fig. 3B,L). In theory, delayed *snail2* activation could be followed by either the later onset of the entire NC specification cascade or by globally defective NC development. To assess definitive NC specification at the end of neural fold stage, we analyzed expression of late NC specification markers, *sox10* and *twist*. While *sox10* was normally activated (Fig. 3C), *twist* expression was severely defective (Fig. 3D) and *hes4*, marking immature premigratory NC was increased (Fig. 3E, Table.S2). In contrast, neural ectoderm, non-neural ectoderm and paraxial mesoderm specification were unaffected (*sox2*, *ep.ker*., *myod*, Fig. 3I-L, Table.S2). Neurulation and neural tube closure were slightly delayed on the injected side, but neural tube eventually closed (Fig. 3.G,I). Importantly, while some neural border markers were expressed normally at the time of neural tube closure (e.g. *msx1*, Fig3.H), others were moderately (*zic1)* to strongly increased (*pax3)* suggesting that the NC cells had partially retained an immature NB-like character, at a developmental stage when they should have been specified into functional premigratory NC (Fig. 3H,I).

**Fig. 3.**
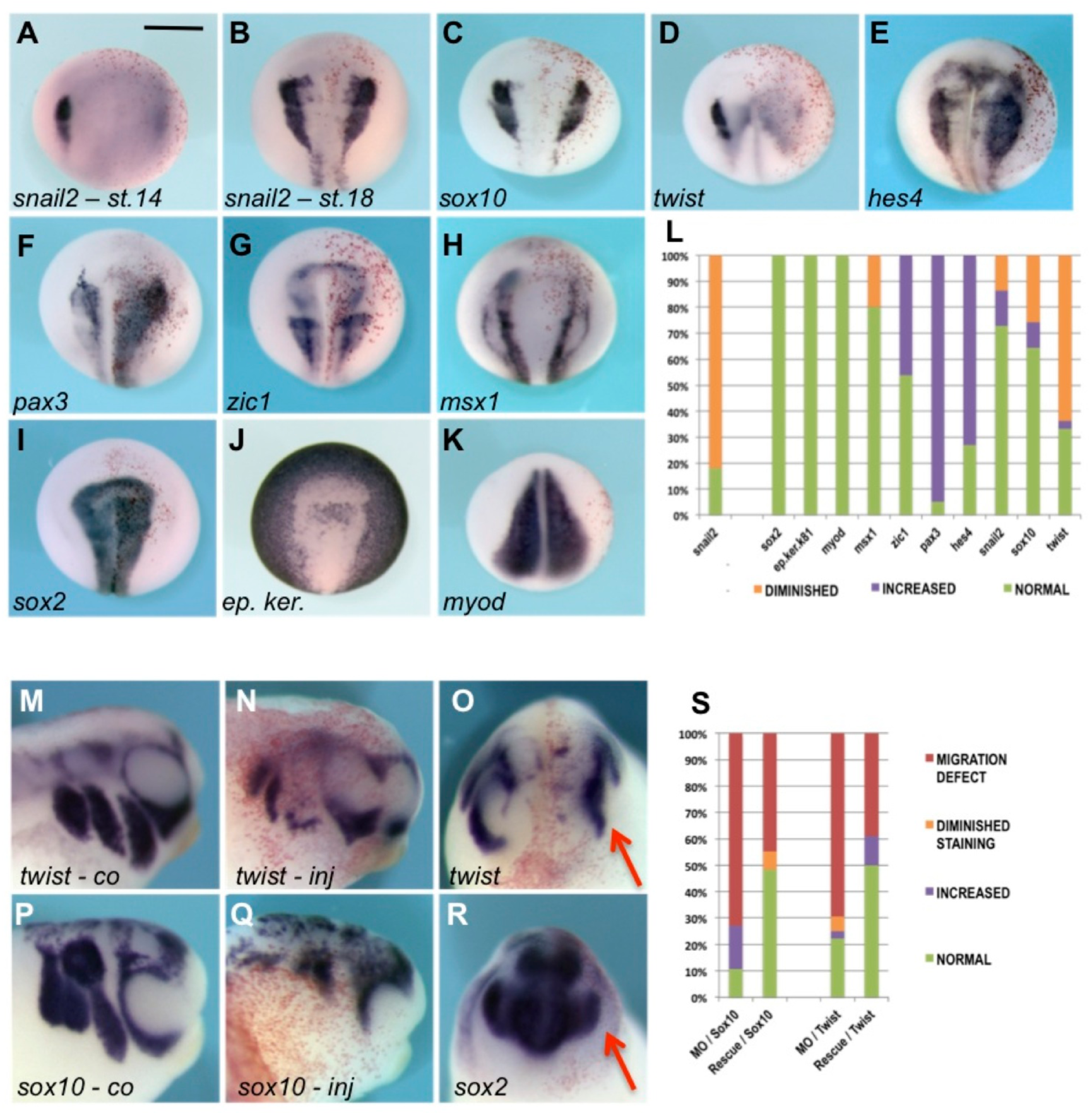
PFKFB4 low-level depletion delays NC early specification, causes retention of NB character, impairs NC late specification and migration. (A) At st.14, *snail2* expression was severely reduced, or abolished, on the injected side (reduced in 78% of embryos, n=42). (B) At st.18, sibling embryos had recovered *snail2* expression (normal in 73% of embryos, n=66). (C-D) While *sox10* expression was mainly unaffected (65%, n=31), *twist* was severely impaired (65%, n=49). (E-H) In contrast, expression of the immature NC marker *hes4* was expanded (73%, n=37), as were some NB markers, either strongly (*pax3*, 100%, n=39) or moderately (*zic1*, 46% n=13). Other NB markers were unperturbed (*msx1*, normal in 80% of embryos, n=10). (I-K) Neural plate (*sox2*, n=13), non-neural ectoderm (*ep.ker*., n=12) and paraxial mesoderm (*myod*, n=4) seemed unaffected. (L) Percent of embryos with each phenotype, details in Table.S2. A-K: dorsal views. (M-R) St.24 tailbud embryos exhibited a severe NC migration defect (M-Q, *sox10*, 50%, n=10; *twist* 56%, n=25). *Sox2* expression appeared grossly unaffected, despite marginal reduction of optic vesicle size (R, n=5). (S) Co-injection of *pfkfb4* mRNA with PFKFBMO rescued both *sox10/twist* alterations of expression and NC migration defects in a significant proportion of the embryos, compared to PFKFB4MO injections: *sox10* and *twist* expression were restored or increased a majority of the embryos (n=29 and n=18 respectively). The injected side (inj) is compared to the control side (co) in side views (M,N,P,Q, anterior to the right) or frontal views (O, R, red arrow on injected side). Scale bar, 500 μm.

After neural tube closure and NC delamination, low-level PFKFB4 depletion resulted in severely defective NC migration, both with diminished expression of migrating NC markers *(twist*, *sox10)*, and decreased migration distance towards the facial areas of the cells expressing these markers (Fig. 3M-S). Although the percent of embryos with abnormal *twist* expression was slightly lower than at the end of neurulation (Fig. 3S-st.24, Fig. 3L-st.18), these results indicated that defects and delay in premigratory NC specification were not compensated at the subsequent stage of NC migration. At this stage, the central nervous system and the optic vesicle were grossly normal (*sox2*, Fig. 3R). When PFKFB4 morphant phenotype specificity was assessed, by co-injecting PFKFB4MO with *pfkfb4* mRNA, the morphant phenotype was rescued during NC specification and migration (Fig. 3S, Fig. S4, and not shown).

Altogether, these results indicated that a moderate loss of PFKFB4 activity resulted in a strong impediment of NC development. The NC territory was established, identified by *sox10* expression and outlined by *msx1* expression (Fig. 3C,H). However, its specification was delayed at neural plate stage (*snail2)*, which was incompletely compensated at late neural fold stage (defective *twist*, increased *hes4*), accompanied by the retention of NB-like character (increased *pax3*) in premigratory NC. This was followed by defective migration of *sox10/twist*-expressing NCC in tailbud stage embryos. As a result, fewer *twist-* or *sox10*-positive cells populated the craniofacial buds. These alterations were sufficient to explain the morphological craniofacial defects observed at st. 45 (Fig. 2). However, using the constitutively active PFKFB4MO, we could not distinguish between an early need for PFKFB4 during NC specification, causing defective NC migration, and a continuous need for PFKFB4 activity during NC specification and migration.

### Inducible PFKFB4 depletion affects neural crest specification and its migration *in vivo*

To understand PFKFB4 function at each step of NC formation, we set up an inducible knockdown strategy, using a UV-cleavable sense morpholino (PFKFB4-PhMO) that blocked *pfkfb4* antisense MO until UV exposure. Although more fragile, we injected albino embryos for UV penetration. The uninjected side controlled for non-specific UV effects. As control, severe PFKFB4MO constitutive depletion led to abolished or severely decreased *twist* expression (in 80% and 17% of embryos, Fig. 4A-C,H). PFKFB4-PhMO/MO co-injections efficiently blocked the severe morphant phenotype (uninduced embryos, u.i.): *twist* was normally expressed in 60% of st.18 embryos. Despite minimal ambient light throughout experiments, we observed some leakage in the PhMO/MO u.i. condition, leading to 22.5% of embryos with severe decrease in *twist* expression and 17.5% with mild defects. However, after UV exposure upon injection (a.i., illumination at the 4-cell stage), *twist* expression decreased significantly on injected side in 73% of embryos (Fig. 4H). The lower efficiency of the PhMO/MO a.i. condition compared to the constitutive PFKFB4MO phenotype, leading to only 26% of embryos with severe *twist* blockade and 47% with mild defects, could be due to a partial *pfkfb4*-*PhMO* cleavage, leading to a hypomorph phenotype. Indeed, the embryos exhibited malformations similar to the ones observed at low doses with constitutive depletion (Fig. 3).

**Fig. 4.**
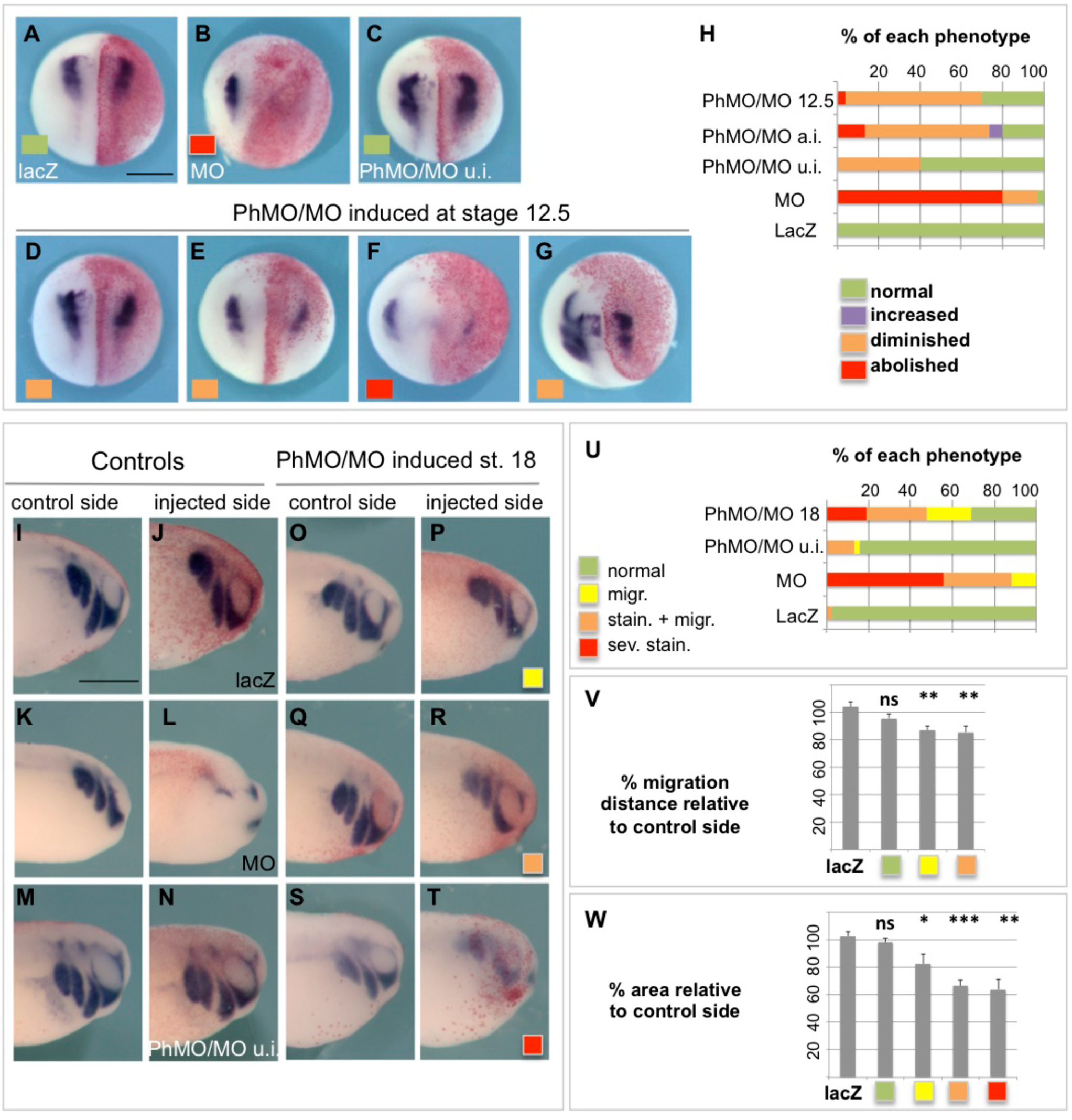
Inducible depletion demonstrates independent PFKFB4 roles in NC specification and migration. We co-injected a UV-cleavable sense MO (PhMO) to block PFKFB4MO until the desired developmental stage and monitored *twist* expression at st18. (A-H) Strategy of phMO validation: *lacZ* control injections (A), unmasked PFKFB4MO, “high” dose (B), PhMO/MO without UV illumination (C, uninduced, u.i); UV illumination at early NC specification st12.5 (D-G). (A) *LacZ* injections did not alter *twist* pattern (100%, n=31). PFKFB4MO severely blocked *twist* expression (abolished (80%) or diminished (17%) n=35). *Twist* decreased in PhMO/MO-injected embryos UV-illuminated immediately after injection (a.i.) (H, 73%, n=15). *Pfkfb4* depletion, activated at st12.5, decreased *twist* expression strongly (47%), or moderately (22.6%) (n=53). This effect persisted until EMT and early NC emigration (G, stage 20). (A-G): dorsal views. (I-W) To deplete *pfkfb4* during EMT/early migration, st. 18 neurulas were UV illuminated and analyzed at tadpole st 24. *Twist* was normal in embryos injected with *lacZ* (I,J; n=41), or with PhMO/MO but not illuminated (M,N; n=38). Cells injected with PFKFB4MO alone died as expected (K,L; n=52). PhMO/MO-injected embryos UV-illuminated at st. 18 exhibited three phenotypes classes (n=42): “migration” (O,P), “staining and migration” (Q,R) and “severe staining” defects (S,T). We compared percent of each phenotype (U), percent of migration distance (V), percent of NC stream area (W, *twist*-expressing area) to the contralateral side. Scale bars = 500 μm. All phenotype differences were statistically significant between illuminated PhMO/MO-injected embryos, and either *lacZ*-injected, or uninduced PhMO/MO-injected embryos (^*^: p-value<0.05; ^**^: p-value<0.01; ^***^: p-value<0.001; ns: non significant).

To study PFKFB4 role during NC specification, embryos were UV-illuminated at the end of gastrulation (st.12.5), and analyzed at late neural fold stage (st.18). Before induction, development appeared normal (*sox2*, *ep.ker*. expression, not shown). PFKFB4 knockdown from st.12.5 resulted in a significant *twist* downregulation (Fig. 4D-H) compared to injected but uninduced embryos. Therefore, PFKFB4 plays a specific role during NC specification, distinct from its early function on ectoderm patterning during gastrulation. Moreover, this effect mirrors the hypomorph phenotype observed with the mild PFKFB4 depletion (Fig. 3).

PFKFB4 knockdown was then induced at late neural fold stage, as NC cells have completed specification, maturation and initiated EMT (st.18). Before illumination, development was mainly normal, despite slightly decreased *twist* expression (u.i., Fig. 4H). In st.24 tadpoles, we analyzed *twist* expression, NC streams shape and migration distance from the dorsal midline. *Twist* was normal in embryos injected with *lacZ* or PhMO/MO uninduced, while the injected cells were lacking (eliminated after cell death) in PFKFB4MO-injected embryos (Fig. 4I-N,U). After st.18 photo-induction, *twist* was significantly altered in 69% of embryos (Fig. 4O-U). For the milder phenotype (“migration”, 21%), tadpoles presented a modestly reduced migration distance (13% reduction), with a robust *twist* expression but smaller NC streams (17% area reduction). This limited effect was reproducible and statistically significant, compared to variations observed in *lacz*-injected embryos or the injected embryos classified as normal (Fig. 4O,P,V,W). The second phenotype (“migration and staining”, 28.5%) presented similarly reduced migration distance (14% reduction) but more intense decrease in NC streams area (34% smaller) and lower *twist* staining intensity (Fig. 4Q,R,V,W). Finally, 19% of the embryos exhibited strongly reduced *twist* expression (“severe staining defect”), with 36% reduced area, the low staining preventing measure of migration distance (Fig. 4S,T,V,W). Hence, depleting PFKFB4 after NC specification, at the time of EMT, affected *twist* expression, the size of NC streams, and NC migration distance.

Together, these results demonstrated a requirement for PFKFB4 function at two successive and distinct steps of NC development: patterning of mature NC during neurulation and NC migration at tailbud stage. Moreover, these effects closely mimicked those observed after low-level PFKFB4 depletion.

### PFKFB4 controls AKT signaling in premigratory NC

PFKFB4 regulation of glycolysis, as seen in adult cells, was not involved in neural crest late specification, since glycolysis blockade during neural fold stage did not alter *twist* expression (Fig. S5, other markers not shown). In order to understand how PFKFB4 may affect NC late specification, we assessed cell signaling parameters and cell proliferation in premigratory NC. In gastrulas, PFKFB4 levels impact PI3K-AKT signaling. Here, to address precisely the level of AKT signaling in developing neural crest progenitors, we dissected out either the neural border territory at the end of neural plate stage, or the premigratory cranial neural crest at the end of neural fold stage, and measured phospho-SER473-AKT levels in control and PFKFB4 hypomorph conditions. AKT signaling was specifically decreased in morphant NC progenitors compared to controls (Fig. 5A)

**Fig. 5.**
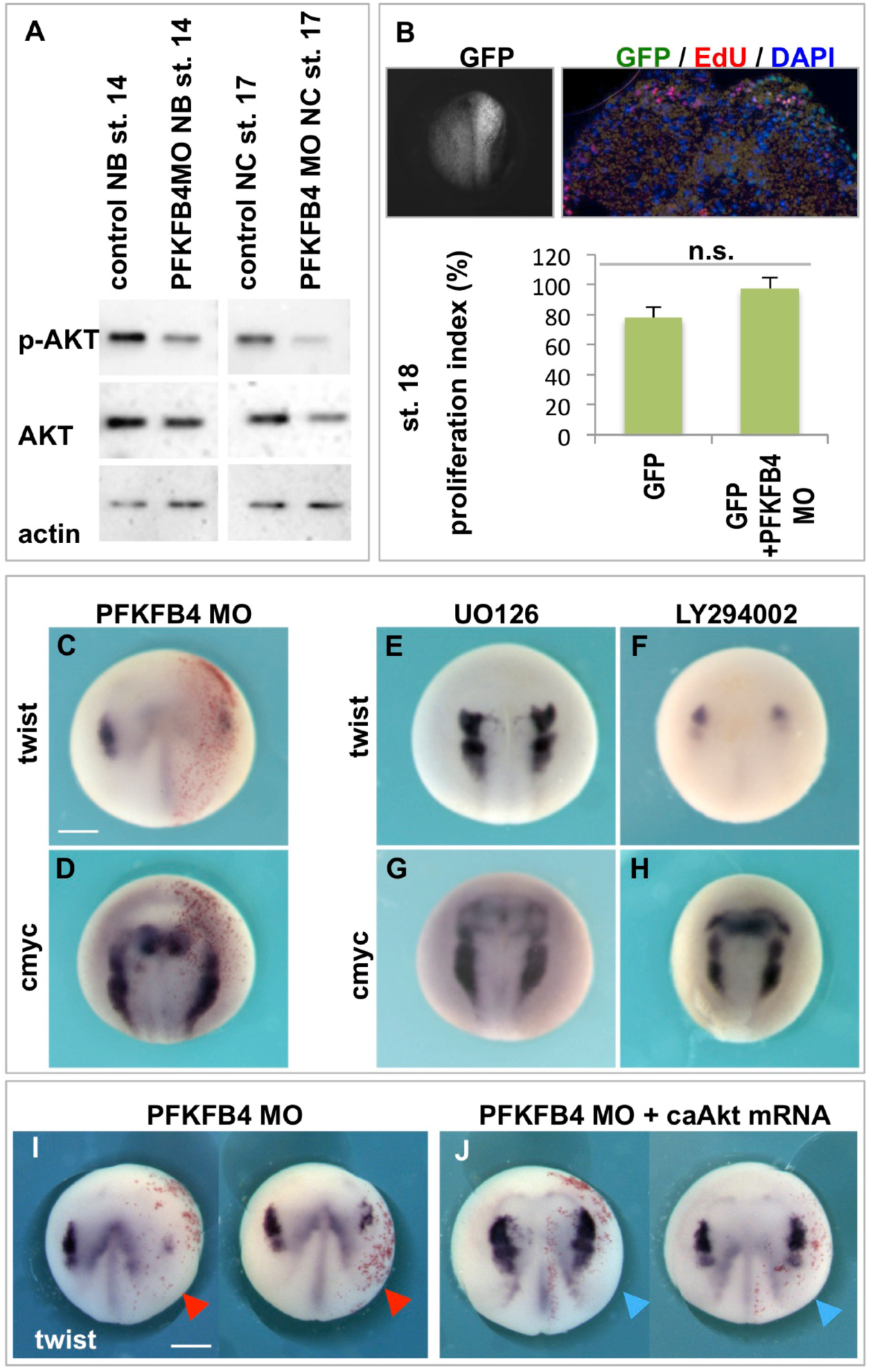
AKT signaling mediates PFKFB4 function on premigratory NC maturation. (A) In embryos, PFKFB4 can regulate AKT signaling in addition to glycolysis rate. Here, in morphants, dissected st. 14 NB or st. 17 NC displayed decreased activated AKT signaling levels. (B) AKT regulates many aspects of cell homeostasis including cell proliferation, cell survival, and cell metabolism. Here, embryos injected with *gfp* and MO in NC area were selected and sectioned: cell counting after EdU incorporation showed that cell proliferation rate was normal after PFKFB4MO injections. (C, D) PFKFB4MO affected late NC specifier *twist* expression, while NC stem cells marker *cmyc* was normally activated. (E-H) Pharmacological treatment during neural fold stage (st. 14 to st. 18) showed that blocking MAPK signaling (E, G) did not affect NC development, while blocking PI3K-AKT signaling (F, H) affected *twist* but not *cmyc*, thus phenocopying PFKFB4MO effect. (I, J) Coinjection of PFKFB4MO with a constitutively AKT (blue arrows) rescued the morphant *twist* phenotype (red arrows): two sibling embryos for each injection are shown. St. 18 *pfkfb4* morphants presented diminished *twist* (69%, n=26). (B) In contrast, siblings co-injected with PFKFB4MO and *caAkt* mRNA had normal *twist* expression in the majority of cases (73% normal or enlarged, n=42). Scale bars = 500 μm.

AKT signaling regulates many aspects of cell homeostasis, including cell proliferation. We next wondered if the rate of cell proliferation was affected in morphant NC, and if the pool of *cmyc*-positive NC stem cells was normal (Bellmeyer et al. 2003). After EdU incorporation, we observed normal cell proliferation on the morphant side (Fig. 5B). *Cmyc* expression was normal on the morphant side, when *twist* was defective in sibling embryos (Fig. 5C,D). Together, these results suggested that the morphant territory generates a rather normal pool of NC stem cells, suggesting that the phenotype relies on genuine patterning defects within a partially specified NC domain.

To assess if diminished AKT signaling was sufficient to affect NC specification and maturation, and to phenocopy the morphant phenotype, we treated embryos with PI3K-AKT inhibitor (LY294002, 40-80μM). As a comparison, we used a MAPK/ERK inhibitor (UO126, 40-80μM) since both AKT and MAPK signaling pathways are often activated downstream of tyrosine kinase receptor signaling. Each treatment was checked for inhibition of p-ERK and p-AKT (on whole embryo lysates, not shown). Embryos were treated exclusively during neural fold stage (from st.14 to st.18). Inhibiting PI3K-AKT signals blocked *twist* but not *cmyc* induction. In contrast ERK inhibitor had no effect on either gene expression (Fig. 5E-H). This result showed that AKT signaling was specifically needed during the last phase of premigratory NC maturation. When AKT function was disrupted, a normal pool of *cmyc*-positive NC stem cells formed but they failed expressing the late NC specifier *twist*.

Finally, when a constitutively active form of AKT was co-injected with PFKFB4MO, *twist* specification defects were rescued, demonstrating that the main cause of PFKFB4 morphant phenotype prior to migration, was due to altered AKT signaling (Fig. 5I-J).

### PFKFB4 depletion affects neural crest EMT and migration in a cell autonomous manner

To understand the defects during NC EMT and migration upon PFKFB4 low-level depletion, we reasoned that morphant NC cells, with delayed or incomplete specification (Fig. 3), might require longer time than wild-type NC to mature and undergo migration. Additionally, interactions between the NC and its *in vivo* environment are essential for migration. Using the more robust pigmented frog embryos, we challenged the ability of morphant NCC to migrate in a wild-type host environment (Fig. 6). We implanted GFP-labeled premigratory morphant NC into a stage-matched wild-type host and followed cell migration *in vivo*. In contrast to the control wild-type grafts which healed within 20 minutes after transplantation, the morphant tissue healed with difficulty: most of the donor tissue failed to adhere to the host tissues, even as long as two hours post grafting (not shown). Moreover, after healing, the vast majority of the grafted morphant cells failed to migrate although occasional cells could be traced in the mandibular arch area (Fig. 6C-D, compare to wild-type cells in A,B). This phenotype was qualitatively similar to the defects observed in the morphant embryos (Fig. 4), either with constitutive or inducible morpholinos. In this transplantation assay, however, the lack of migration was observed in a higher number of cases, possibly because of the additional healing defect revealed by the assay. This challenge revealed that PFKFB4 morphant cells displayed altered healing and survival ability upon EMT and migration stage.

**Fig. 6.**
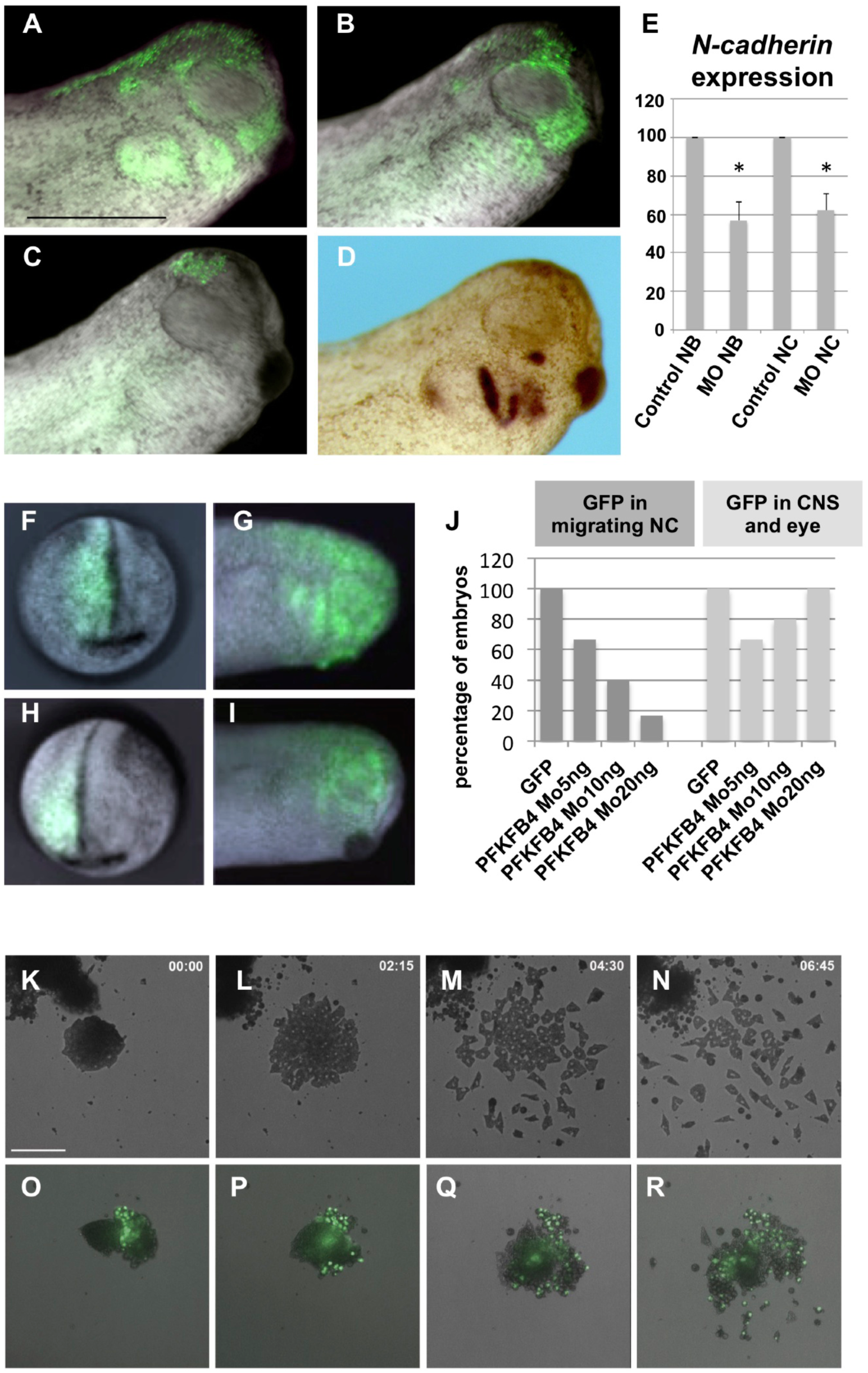
PFKFB4 morphant neural crest progenitors fail to undergo EMT and migration. (A, B) When transplanted into a wild-type host embryo, GFP-labeled wild-type NC efficiently migrates and populates the host branchial arches, whichever the size of the grafted tissue (A) large graft, (B) small graft (20/24 cases, n=24). (C, D) In contrast, PFKFB4 morphant NC exhibited defective healing, resulting in small-size grafts, which yielded few, if any, migratory NCC into the host craniofacial area (n=23, occasional migration in 9/23 grafted embryos). (C) Small graft without migratory cells and (D) small graft with few cells reaching the branchial arches, *gfp* staining by WISH to enhance individual cell visualisation. (E) *N*-*cadherin* expression (RTqPCR on dissected explants), either prior to EMT (NB, st. 14), or in premigratory NC (st. 17). (F-J) Wild-type or morphant cells were lineage traced *in vivo*, on embryos injected in the prospective neural fold unilaterally (F, H). (G-J) At tailbud stage, control cells (G) efficiently populated branchial arches and were also found in brain and eye (J). In contrast, morphant cells failed to populate branchial arches (I) in a dose-dependent manner (J), but were normally found in the brain and eye (J). (K-R) NB were dissected prior to EMT (st. 14), plated onto fibronectin. Cell behavior was followed by time-lapse videomicroscopy. Wild-type NC (K-N) adheres efficiently, undergoes EMT (L), cell scattering and migration (M, N). Morphant NC (O-R) presented poor adherence, delayed (P) and inefficient EMT (Q), and few emigrating cells (R). Scale bar = 160 μm.

We further analyzed the fate of the morphant cells *in vivo* with another strategy. In order to avoid interference with altered healing properties, we traced the injected cells with *gfp* mRNA co-injected with PFKFB4MO, by targeting dorsal-animal blastomere D1.2 at stage 8 to 16 cells, which mostly forms dorsal neural tube and NC progenitors (Moody et al., 1987). We followed the fate of NC progenitors, by selecting embryos with GFP-positive premigratory NC at the end of neurulation (st.18, Fig. 6F-H). In these conditions, without further experimental manipulation of the injected cells, we find that the control cells undergo EMT, migrate and populate efficiently craniofacial areas as expected, while morphant cells fail to do so in a dose dependent manner. In contrast, the morphant cells contribute to brain and eye as efficiently as control cells (Fig. 6G,I,J). We concluded that the defective NC specification and maturation, observed at st.18, was not compensated later on, even when morphant cells face a wild-type environment, and that a lower PFKFB4 activity resulted in a cell-autonomous alteration of NC ability to undergo EMT and migration *in vivo*.

To understand the cellular and molecular basis of morphant NC phenotype, we tested adherence, EMT and migration on fibronectin *in vitro* (Fig. 6K-R). This assay was done using earlier tissue than in previous publications, i.e using neural border at st. 14 instead of premigratory NC st. 17-18, which has already started EMT. Doing so, this assay allows visualizing separately adherence a few minutes after plating, then EMT about 2 hours after plating, and finally cell migration on the fibronectin substrate starting 3,5-4hours after plating (Fig. 6K-N). Morphant neural folds were thus dissected out at the end of neural plate stage (st.14). They failed to adhere efficiently on fibronectin, did not undergo EMT (O-P) and poorly migrated (Q-R), while wild-type neural folds adhered, underwent a very clear EMT (K-L), dispersed and actively migrated (Fig. 6M-N)). A key parameter for NC EMT is the up-regulation of *n*-*cadherin* expression prior to EMT (Theveneau & Mayor 2012). We found that *n*-*cadherin* expression levels were significantly lower in morphant NB (st.14) and NC (st.18) compared to stage-matched controls (Fig. 6E).

We concluded that lower PFKFB4 activity resulted in premigratory NC with defective *n*-*cadherin* levels. These cells were unable to undergo EMT on a fibronectin substratum *in vitro*, exhibited poor ability to adhere, heal, or migrate upon grafting in wild-type host environment *in vivo*. The morphant cells remained integrated into adjacent tissues such as central nervous system or eye. These altered capacities were sufficient to explain the late phenotype in tadpoles, with underdeveloped craniofacial structures.

### At tailbud stage, PFKFB4 depletion alters both glycolysis and AKT signaling, which together impact NC migration

PFKFB4 depletion decreased glycolysis in embryos (Fig. S5). To understand PFKFB4 depletion phenotype during NC migration, we blocked either glycolysis or AKT signaling from EMT stage (st.18) to migration into branchial buds (st.24) (Fig. 7, S6). Glycolysis was blocked using 2-deoxyglucose (2DG), a non-hydrolysable glucose. The PI3K inhibitor LY294002 was used at various doses to block PI3K-AKT signaling. Efficiency of each treatment was monitored (Fig. S5). We found that both glycolysis and AKT phosphorylation blockade from st.18 to st.24, resulted into severe disruption of NCC migration, (*twist*, *sox9*, *sox10*, Fig. 7, Fig. S6). The effects of LY294002 were dose-dependent (not shown). While glycolysis blockade seemed to mainly affect NC streams morphology, AKT inhibition also affected general embryo development (not shown). In order to test for delayed NCC migration and for general treatment toxicity, embryos were transferred in drug-free medium from tailbud st.24 until late tadpole st.45. In these conditions, potential delays would be compensated, while general toxicity would lead to embryo death. Here, embryo survival was normal. St.45 craniofacial morphology was analyzed. Relative to body size, heads were smaller and visceral cartilage elements were severely underdeveloped as observed upon PFKFB4 depletion, especially after AKT signaling blockade. Interestingly, combining both glycolysis and AKT blockade led to stronger craniofacial phenotype, suggesting cooperation of the two processes (Fig. S6). We concluded that blocking AKT signaling or, with less efficiency, blocking glycolysis, phenocopied PFKFB4 morphant phenotype during NC migration. Both AKT signaling and glycolysis were thus needed for a large number of NCC to populate the branchial arches and head structures.

**Fig. 7.**
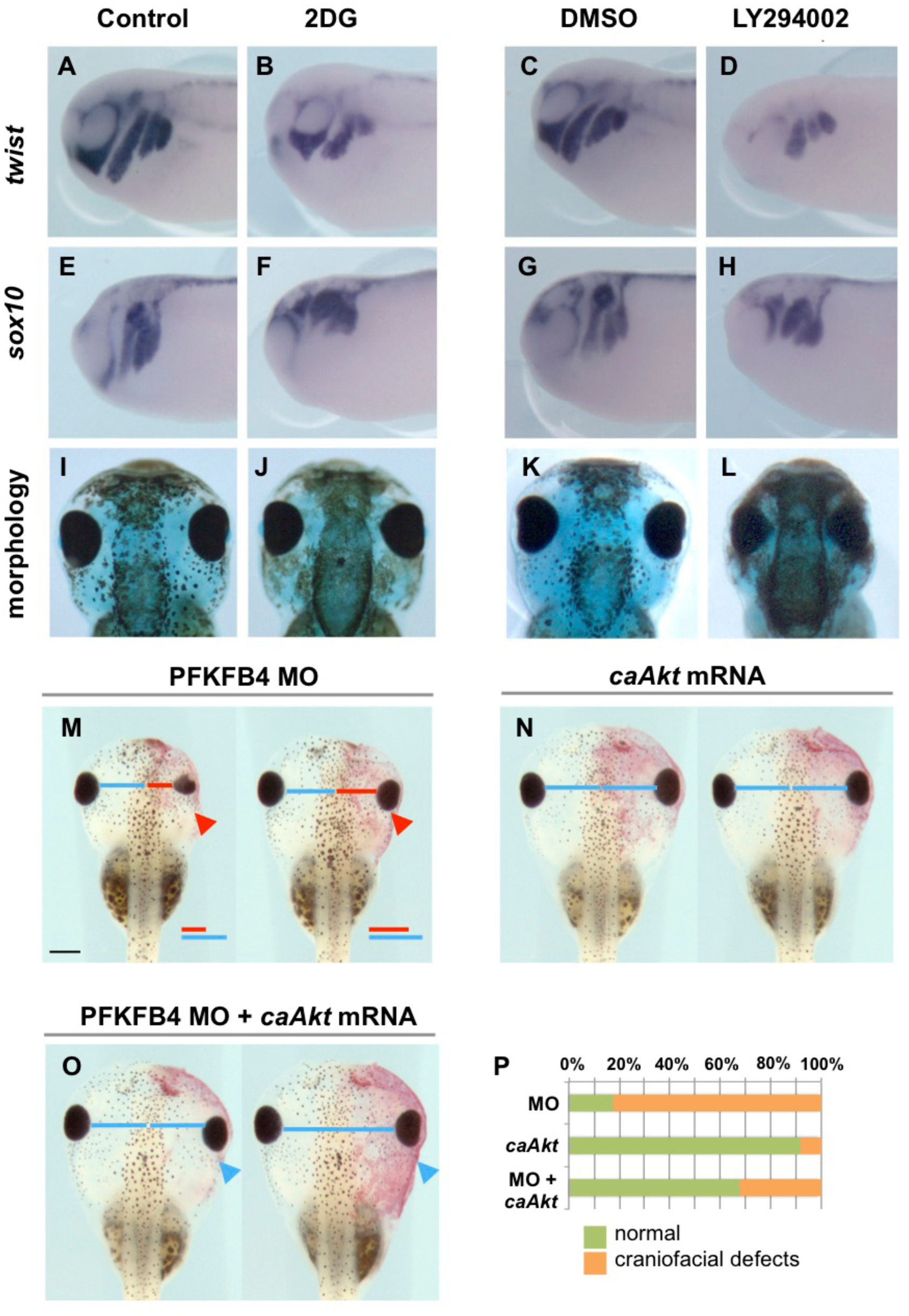
Glycolysis and PI3K-AKT signaling impact NC migration similarly to PFKFB4 low-level depletion, and activating AKT signaling rescues PFKFB4 downregulation. (A-H) When glycolysis (2DG) or PI3K-AKT pathway (LY294002) were blocked during EMT and migration (st. 18 to st. 24) both treatments severely affected NC migration at st. 24 (*twist*, n=16 (2DG), n=14 (LY); s*ox10* (n= 11 (2DG), n=12 (LY). (I-L) At st. 45, tadpoles treated during NC migration then grown in control medium, exhibited general head morphology defects, including eye defects and reduced branchial cartilages. (I, K) sibling controls, n=30; (J) 2DG, n=13; (L) LY294002, n=19. (A-H) side views, (I-L) dorsal view. (M) At tadpole st. 45, morphant sides were severely affected (P, 83% embryos with craniofacial reduction, n=23), while activation of *Akt* signaling (N) did not affect overall craniofacial morphogenesis (P, 92% symmetrical, n=59). Tadpoles co-injected with PFKFB4MO and *caAkt* (O) were largely rescued, with 66% of embryos with injected side symmetrical to contralateral side (P, n=63). (M) Red bar: eye distance from the midline (morphant side). Blue bar: control distance. Both bars were aligned for comparison. (N) On both sides, the same blue bar measures eye distance from the midline. Scale bar = 500 μm.

Since AKT blockade seemed most effective in mimicking PFKFB4 depletion, we attempted to rescue the craniofacial phenotype by co-injecting *caAkt* mRNA with PFKFB4MO. By st.45-46, we observed that constitutive activation of AKT signaling alone did not affect craniofacial development (Fig. 7N,P). While morphant injected sides were severely reduced (Fig. 7M,P), most tadpoles co-injected with PFKFB4MO and *caAkt* were symmetrical (Fig. 7O,P). This experiment demonstrated that defects of NC craniofacial derivatives observed upon PFKFB4 depletion were largely due to loss of AKT phosphorylation and were efficiently compensated by restoring active AKT signaling.

### The NC-GRN regulators control *pfkfb4* activation at the neural-non-neural border

Finally, we analyzed *pfkfb4* upregulation at the edges of the neural plate. We asked if the NC-GRN actors actively controlled *pfkfb4* expression, as they do for *snail2*, either during early *pfkfb4* activation in the end of neural plate stage, or in premigratory NC. We used MO-mediated depletions with well-established morpholinos, dominant-negative constructs (Fig. 8) or gain-of-function experiments (Fig. S7). First, the secreted NC inducers, including WNT and FGF8 signals as well as modulating BMP signaling, affected similarly *snail2* and *pfkfb4* (Fig. 8A-L, Fig. S7). Secondly, we found that the key transcription factors acting at the NB to specify NC, PAX3 and TFAP2a, were also essential to establish *snail2* and *pfkfb4* (Fig. 8M-T, Fig. S7). In addition, the NC specifier SOX9 was needed for expression of both genes at the NB and in premigratory NC (Fig. 8U-X). All these results indicated that increased *pfkfb4* expression in the NC progenitors is actively promoted by the NC-GRN regulators, as for more classical partners of the network (e.g. *snail2*). Thus, our results show that PFKFB4 key function, which mainly relies on ensuring proper AKT signaling in NC progenitors, is encoded within the NC-GRN. PFKFB4 could be a major intermediate actor in NC control by WNT-FGF-BMP signals.

**Fig. 8.**
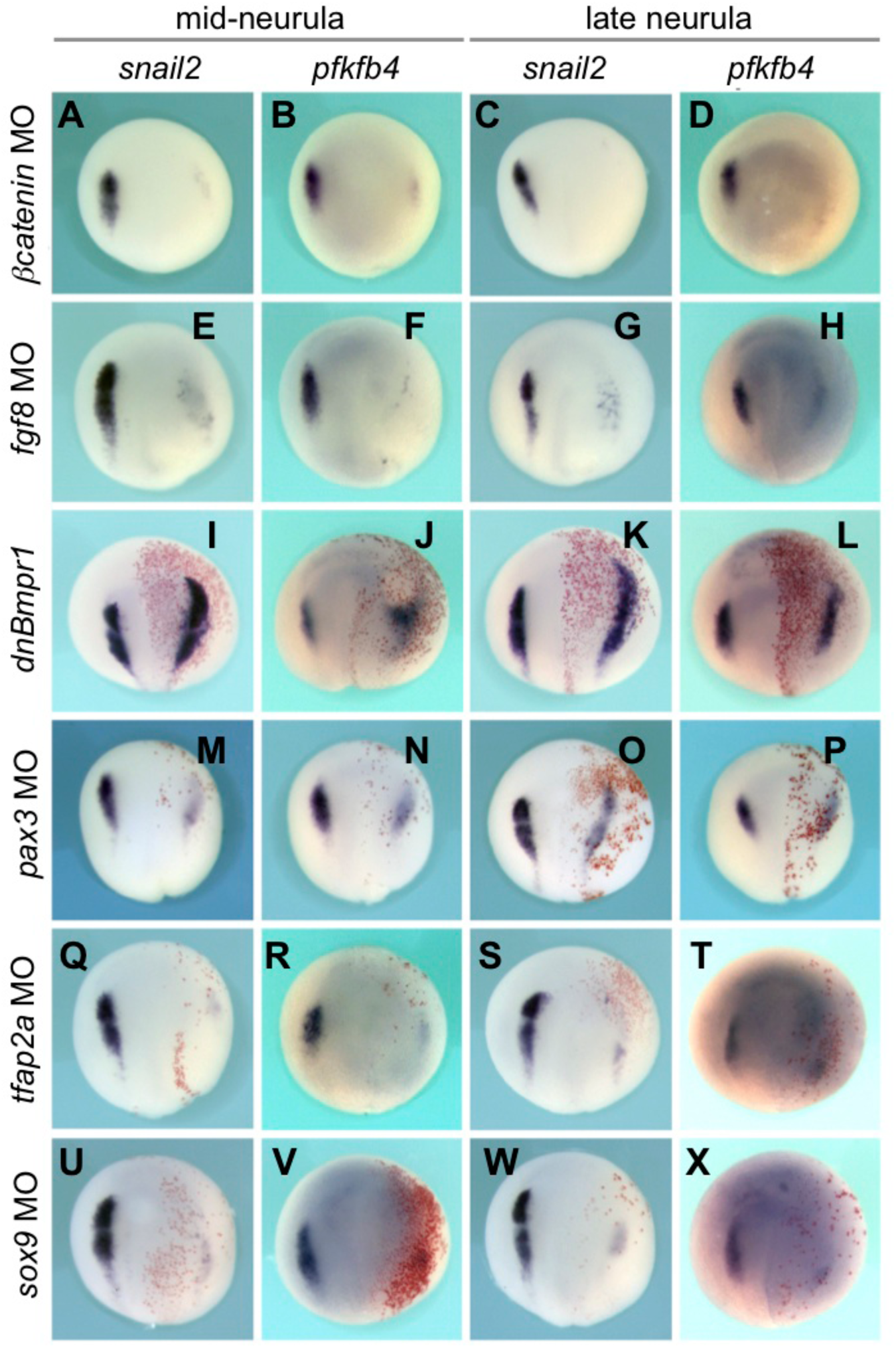
*Pfkfb4* activation during neural fold stage is regulated by the neural crest GRN. (A-Y) Unilateral depletion of members of the NC-GRN modulated *pfkfb4* activation at the NB (st.14) and NC (st.18). (A-H) Blockade of WNT and FGF signaling strongly decreased *snail2* and *pfkfb4* expression. (I-L). Conversely, BMP signaling downregulation upregulated *snail2* and *pfkfb4* expression, laterally to the NB. (M-T) Knockdown of the NC specifiers PAX3 and TFAP2a resulted in loss of *snail2* and *pfkfb4*. (U-Y) Knockdown of the NC specifier SOX9 depleted *snail2* and *pfkfb4*. Dorsal views, injected side on the right. Detailed scoring: see Table S2. Scale bar = 500μm.

## DISCUSSION

In this study, we have evidenced that PFKFB4 activity is required continuously during several key steps of NC formation: during premigratory NC maturation at neural fold stage, and during NC migration towards the craniofacial buds at tailbud stage. This activity is critical as PFKFB4 depletion led to hypomorphic craniofacial structures. We showed that the major transcription factors and signaling pathways that control the NC-GRN, control *pfkfb4* activation in NC progenitors during neural fold stage, i.e. during the phase of NC maturation. In turn, PFKFB4 controls AKT signaling activity in NC, which is essential for NC maturation and migration. PFKFB4 is a known glycolysis regulator, and its depletion in neurulas affected glycolysis. However, at neural fold stage, glycolysis is not required for NC patterning. In contrast, at tailbud stage, both AKT signaling and glycolysis are important parameters to ensure NC migration. Finally, restoring AKT signaling compensates PFKFB4 depletion and results in normal craniofacial development. This indicates that PFKFB4 critical action on NC development is mainly mediated by its action on AKT signaling (Fig. 9).

**Fig. 9.**
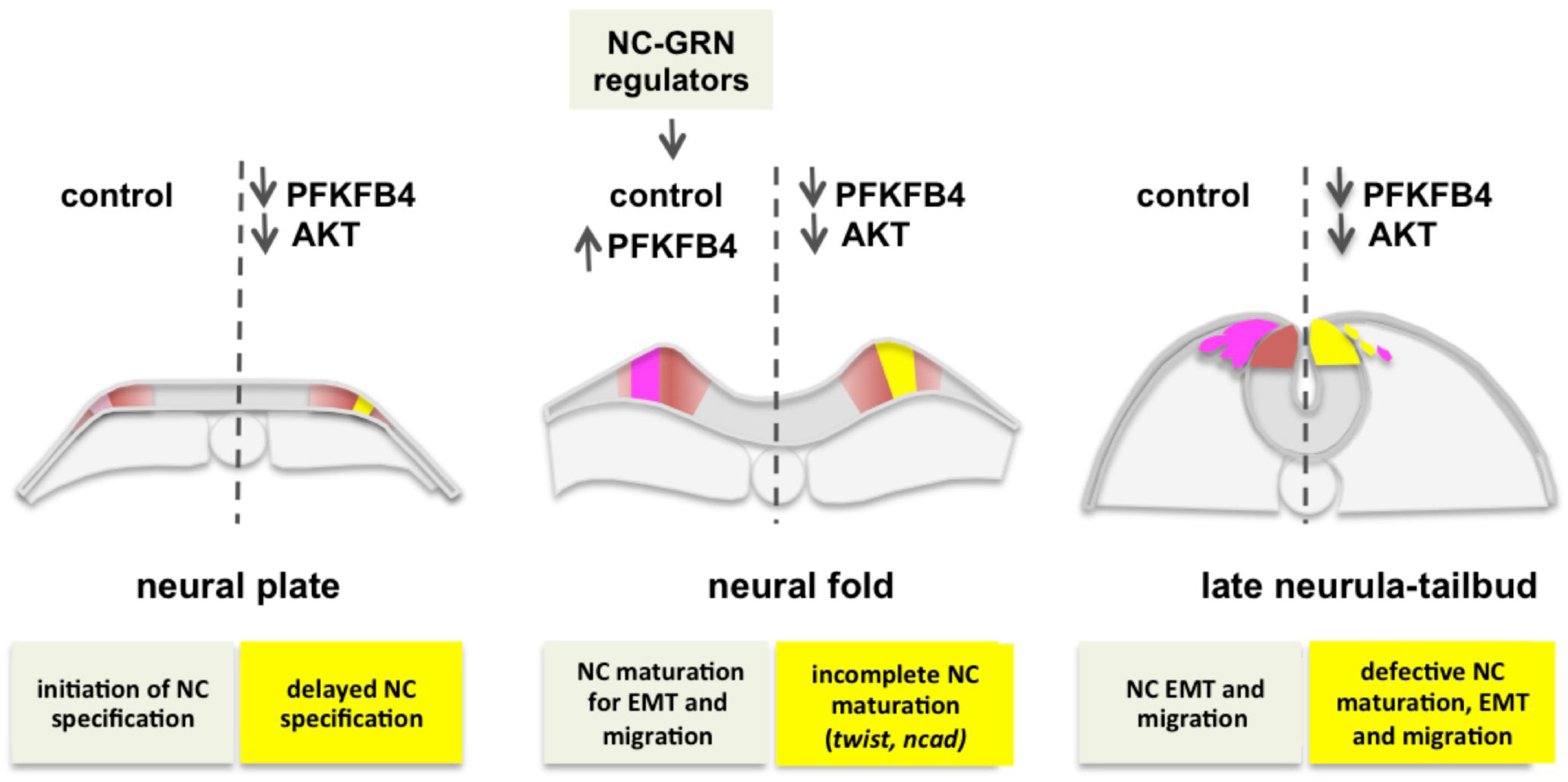
Model of PFKFB4-controlled AKT signaling in NC maturation, EMT and migration. During neurulation, in control conditions (left side), NC induction is initiated at the neural plate stage with weak *snail2* expression. This initial induction is then strengthened during neural fold stage, under the action of the NC-GRN which, in particular, activates *pfkfb4* expression in NC progenitors. This second phase allows acquisition of cellular ability to undergo EMT and migration upon neural tube closure. When AKT signaling is defective (right side), either after PFKFB4 low-level depletion, or using pharmacological inhibitors, delays in NC specification cascade occur, incomplete maturation is observed, and NC fails to undergo EMT. When PFKFB4 and AKT functions are prevented after EMT, fewer NC cell migrate. As a result, reduced craniofacial skeletal elements form. This series of results shows that the steps of NC early development rely upon a continuous and elevated AKT signaling level, sustained by PFKFB4, itself triggered by the NC-GRN.

The enzyme PFKFB4 is an integral partner of the NC-GRN. While it is ubiquitously expressed in the dorsal ectoderm at gastrulation stage (Pegoraro et al., 2015), we found that, during neurulation, *pfkfb4* expression is upregulated in immature NC progenitors, starting at neural fold stage. *Pfkfb4* is further expressed in the NC initiating their migration (Fig. 1. We have shown that the major NC-GRN regulators, including secreted inducers, NB specifiers, and NC specifiers, control *pfkfb4* expression in the neural folds (Fig. 8). Moreover, depletion of PFKFB4 did not prevent NB specifiers expression but led to delayed or decreased expression of early (*snail2)* and late *(twist)* NC specifiers (Fig. 3). According to the hierarchical model of the NC-GRN, this indicated that PFKFB4, while not a transcription factor, acts as a NC specifier (Betancur et al. 2010; Milet & Monsoro-Burq 2012). The maintenance of *pfkfb4* expression in NC could further involve NC-GRN actors. We have evidenced that PFKFB4 was needed reiteratively during NC specification and migration (Figs.4,6). Likewise, WNT and TFAP2a function are also needed at successive steps of NC formation (LaBonne & Bronner-Fraser 1998; Monsoro-Burq et al. 2005; Simões-Costa et al. 2015; Luo et al. 2003; de Crozé et al. 2011). Both WNT signals and TFAP2a activate *pfkfb4* expression during NC specification, suggesting that they could potentially sustain its expression also during later NC development.

Our data suggest that the main function of PFKFB4 upregulation in premigratory and migratory NC is to ensure optimal AKT signaling levels in these cells (Fig. 5, 7). We also show that blocking AKT signaling in temporally controlled conditions, affects NC specification and migration independently (Fig. 5, 7). In mouse embryos, a recent study also demonstrated importance of AKT signaling for NC migration: *Specc1l* mutants or knockdowns exhibit defective neural tube closure and cranial NC migration, with decreased phospho-AKT levels. In this context also, activating PI3K-AKT signaling rescued the phenotype (Wilson et al. 2016). Together these data suggest that the acute need for optimal AKT signaling during NC migration might be conserved in amniote and non-amniote vertebrates.

The regulation of NC early development by PFKFB4 and AKT was essential to shape the embryonic head. We propose that it ensures that numerous NCC populated the branchial arches, and differentiated into cartilage elements with appropriate size rather than impacting the morphogenesis of these elements (Fig. 2, 7). When inducible depletion was performed during NC migration only, cells eventually migrated almost down to the target tissues, but in smaller numbers (Fig. 4). Neural tube closure and craniofacial defects are an acute societal issue for human health. Hypomorphic craniofacial structures, especially for the jaw, are observed in many human syndromes, and not yet linked to specific mutations (reviewed in Heike et al. 1993). Our study shows that impaired AKT signaling, specifically in the NC progenitors by means of PFKFB4 depletion, creates reduced jaw and branchial arches structures. This could happen because defects arose at time of NC specification, or during its migration. Our results thus show that even modest reduction of PFKFB4 levels, or of AKT signaling, when applied during developmental periods corresponding to critical steps of NC development, results into severe craniofacial defects. This study highlights the importance of a strict temporal schedule during NC developmental cascade. This schedule culminates with the EMT and migration onset of mature NC cells in register with neural tube closure. Optimal AKT signaling, regulated by high PFKFB4 levels, ensures that no delay in this developmental cascade occurs, that immature progenitor (NB-like) characters are not retained, and that the ultimate molecular switches (*twist and n*-*cadherin* expression) are activated in a timely manner, allowing NC to undergo EMT upon neural tube closure. Later on, PFKFB4 and AKT signaling further optimize the efficiency of NC cell migration towards the branchial arches. Similarly, AKT signalling was recently shown to up-regulate *n*-*cadherin* and control NC migration downstream of PDGF-A/PDGFRa (Bahm et al., 2017). This suggests that various cell inputs use AKT signaling as a hub to control NC formation.

During NC migration, our results outlined a potential link between PFKFB4, AKT and glycolysis, because AKT activation rescued the global *pfkfb4* morphant phenotype (lower AKT signaling and lower glycolysis rate, Fig. 5, 7). We conclude that AKT compensates all aspects of PFKFB4 depletion, directly or indirectly. In adult cancer cells and stem cells, PI3K/AKT signaling regulates glycolysis, which controls tumor cell motility (Han et al. 2013; Ito & Suda 2014; Courtnay et al. 2015). This mechanism could be involved in NC, as forced AKT signaling rescues PFKFB4 phenotype. Comparison with cancer cells further highlight potential common mechanisms. In tumors, glycolysis enzymes are multifunctional regulators, which impact on cell physiology in addition to their role in glycolysis. For example, Alpha-enolase regulates PI3K-AKT signaling and downstream cell EMT regulators such as *snail1* and *n*-*cadherin* in lung cancer (Fu et al. 2015). PFKFB3 controls PI3K-AKT in human osteoarthritis cartilage, in adipocytes and in cancer cells (Trefely et al. 2015; Qu et al. 2016). It has been proposed recently, that increased aerobic glycolysis in tumor cells (Warburg effect) promotes cell survival and cell cycle by stimulating IGF/AKT signaling, in addition to its role in cellular energy and metabolic intermediate production (Trefely et al. 2015). Our findings highlight a novel embryonic process, essential for NC EMT and migration, which parallels some mechanisms of cancer progression. It was shown that migrating NC cells displayed increased resistance to multiple cell stresses (Vega et al. 2004), ensuring their optimal long distance travel throughout various embryonic environments. Altogether, our findings show how these properties are acquired and sustained, during and after premigratory NC maturation, and how they are encoded in the NC-GRN, using unconventional function of a glycolysis regulator to control AKT signaling.

## MATERIALS AND METHODS

All morpholinos, plasmids and reagents are described in Table S1.

### Embryos, microinjection, NB culture and NC grafts

Pigmented and albino *Xenopus laevis* embryos were obtained and staged using standard procedures (Sive et al. 2010) (Nieuwkoop & Faber 1994). European and National Regulation for the Protection of Vertebrate Animals used for Experimental and other Scientific Purposes were strictly applied (licence #C91-471-108, *Direction Départementale de Protection de la Population*, *Courcouronnes*, France). MOs or mRNAs were co-injected with *NLS*-*lacZ* mRNA or *histone2B*-*gfp* mRNA for lineage tracing. Bgalactosidase activity, revealed in red, marked the injected side. Embryos were injected unilaterally for WISH/morphology or on both sides (for western-blot and RTqPCR).NB and NC culture/grafting experiments were described in (Milet & Monsoro-Burq 2014).

### Plasmids and morpholinos

mRNAs were obtained *in vitro* (mMessage mMachine SP6 or T7 kits, Ambion). UV-cleavable photomorpholino (PhMO) experiments were performed on albino embryos (molecular ratio of sensePhMO/splice-blockingMO=1.25). PhMO cleavage was induced by 15 min UV exposure using a HBO 103W/S source.

### Pharmacological treatments

Glycolysis, MAPK and PI3-Akt signaling were inhibited using 2-deoxyglucose (Sigma), UO126 or LY294002 (Sigma) respectively, on batches of 50 sibling embryos. Lactate concentration was measured on 20-40 embryos (L-Lactate Kit, Abcam).

### Cell proliferation and cell death assays

Cell proliferation was assessed based on EdU incorporation for 2 hours (1mM EdU injected ventrally into st. 18 embryos, Molecular Probes). Either *gfp* mRNA alone, or PFKFB4MO and *gfp* mRNA were injected unilaterally. Serial paraffin sections were immunostained. A minimum of 130 cells in the territory corresponding to the neural crest was counted on the GFP-injected side, compared to the contralateral side. Cell death was assessed by activated caspase 3 immunostaining *in toto*.

### Cartilage staining

Embryos were stained with 0.05% Alcian blue, destained in ethanol, rehydrated and cleared with 4% KOH then in graded glycerol solutions. Cartilages were manually dissected out.

### Whole-mount in situ hybridization (WISH) and sectioning

We used a procedure optimized for NC (Monsoro-Burq 2007). For sectioning, embryos were embedded in gelatin-albumin. 30 μM thick vibratome transverse sections were cut.

### Western blotting

Lysates (5-10 whole embryos, 10-15 NBs, NCs or dorsal explants) were prepared using phosphatase (PhosSTOP, Roche) and protease-inhibitors (Sigma) and analysed by standard western blotting.

### RT–qPCR

Total RNA was extracted from 3-5 whole embryos or 5 NB, NC or dorsal explants (Sive et al, 2000). RT–qPCR was performed according to standard procedures using MIQE recommendations. Results were normalized against the reference genes *odc* and *ef1a*. Primers used to test PFKFB4MO efficiency span the exon1-exon2 region, i.e. include the exon1-intron1 junction targeted by the morpholino.

### Statistical analysis, imaging and image processing

All experiments were performed at least three times independently (except duplicates for Figs. 5A). The most frequent phenotypes are shown. Graphs indicate the mean percentage of embryos for a given phenotype. NC stream area and NC distance of migration was measured (Fig. 4, ImageJ software). Error bars = SEM. Student’s t-test was used (p-value ≤0.05). Images were processed with standard calibration of RGB levels (Photoshop, Gimp).

## ACKNOWLEDGEMENTS

The authors are grateful to all members of the Monsoro-Burq team and to S. Saule for interactive discussions and R. Harland for gift of plasmids. We specially thank C. Milet, A. Valluet and A. Dolly for initiating the Akt analysis, C. Pouponnot and A. Eychène for IP experiment. We thank the PICT-IBiSA imaging facility for technical advice and E. Belloir and C.Alberti for *Xenopus* husbandry.

## AUTHOR CONTRIBUTIONS

A.L.F, F.M, C.B, P.P, C.P, M.S. and A.H.M.B designed, conducted and analyzed the experiments. A.L.F, F.M, C.B, P.P, M.S. and A.H.M.B have discussed the results and written the manuscript.

---

**Table S1.**
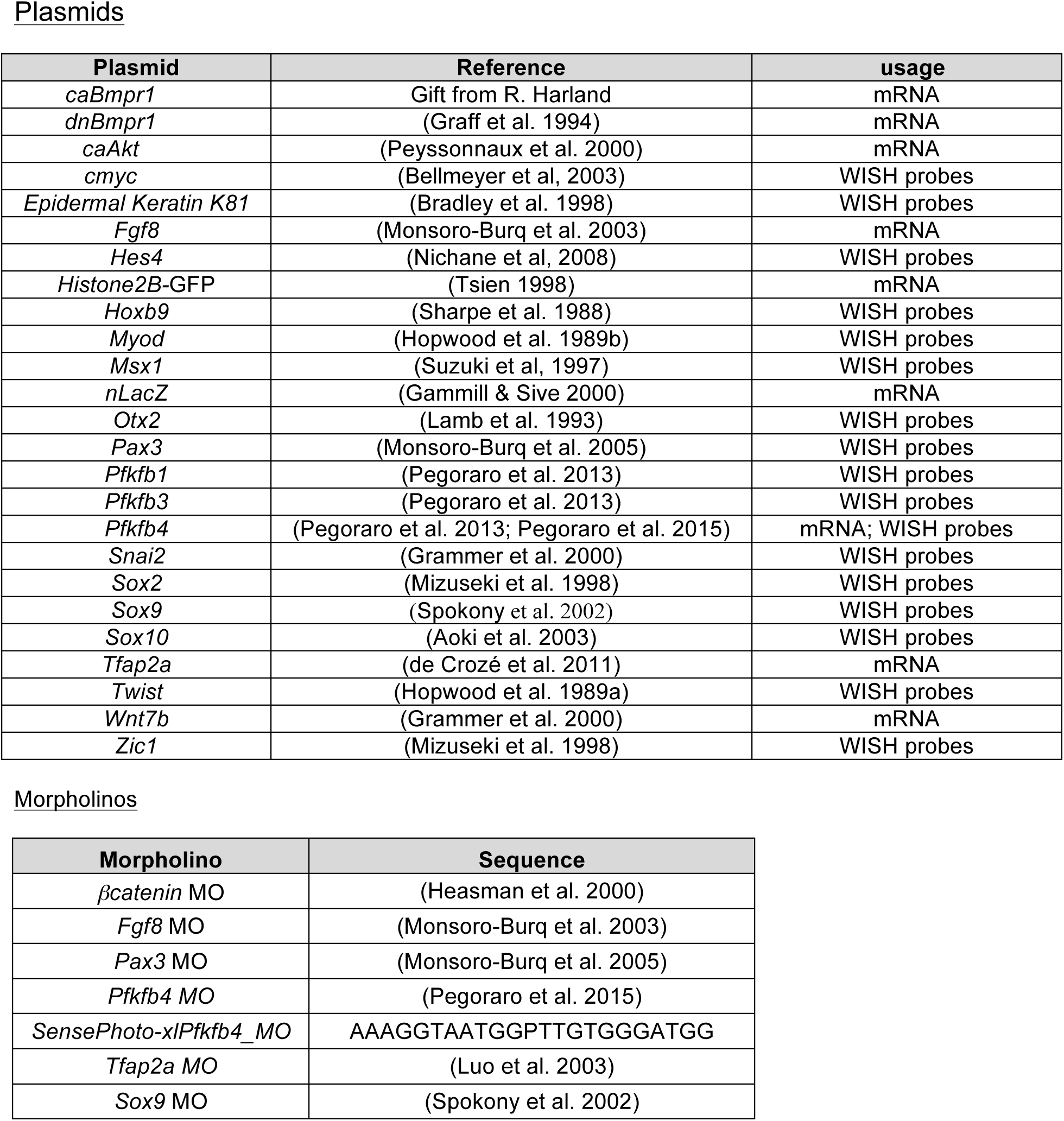
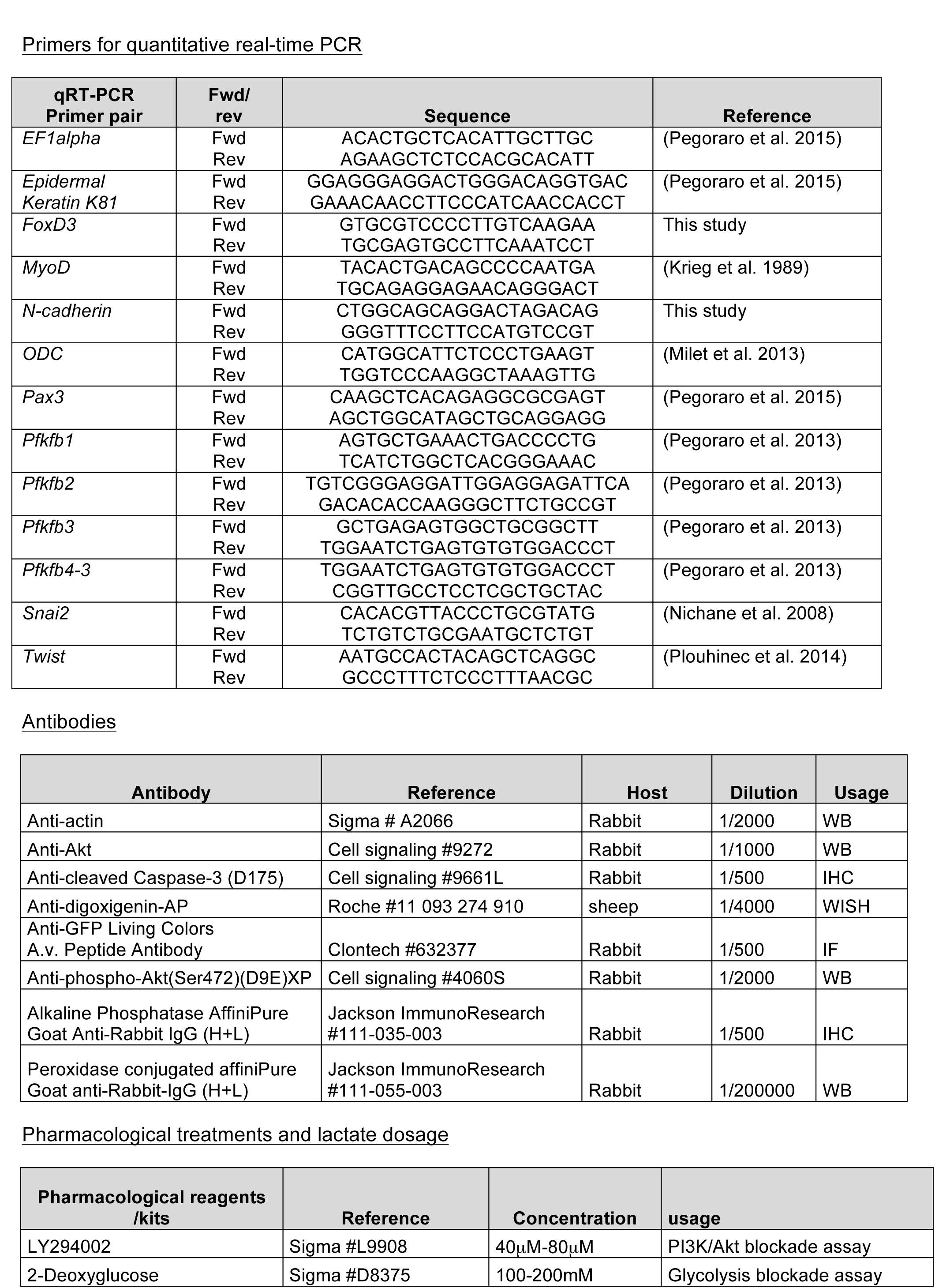
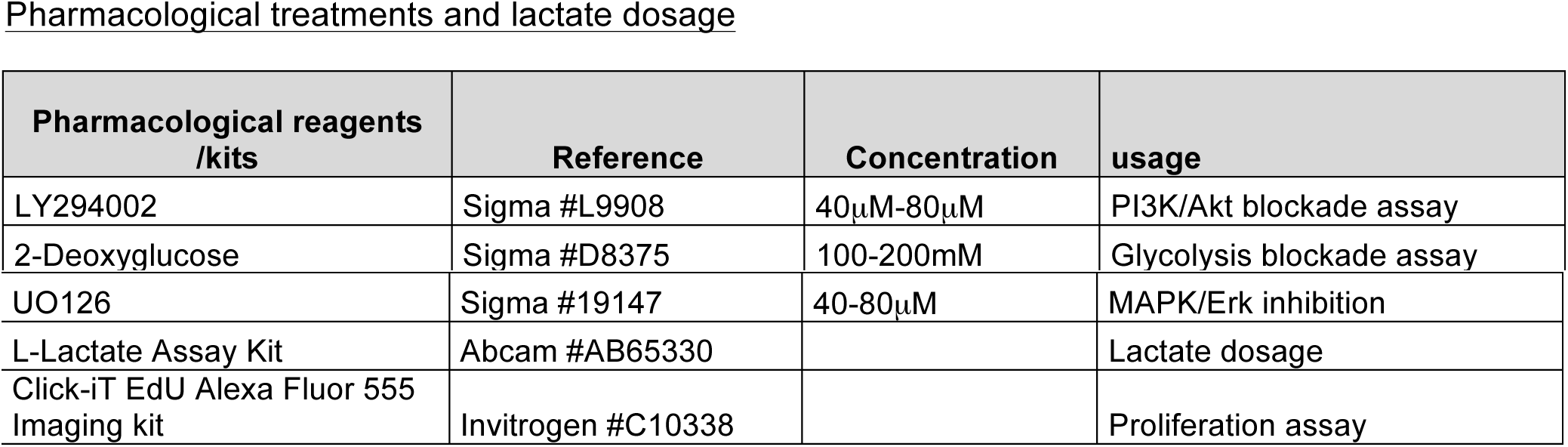
Reagents used in this study

## SUPPLEMENTARY FIGURES LEGENDS

**Fig. S1.**
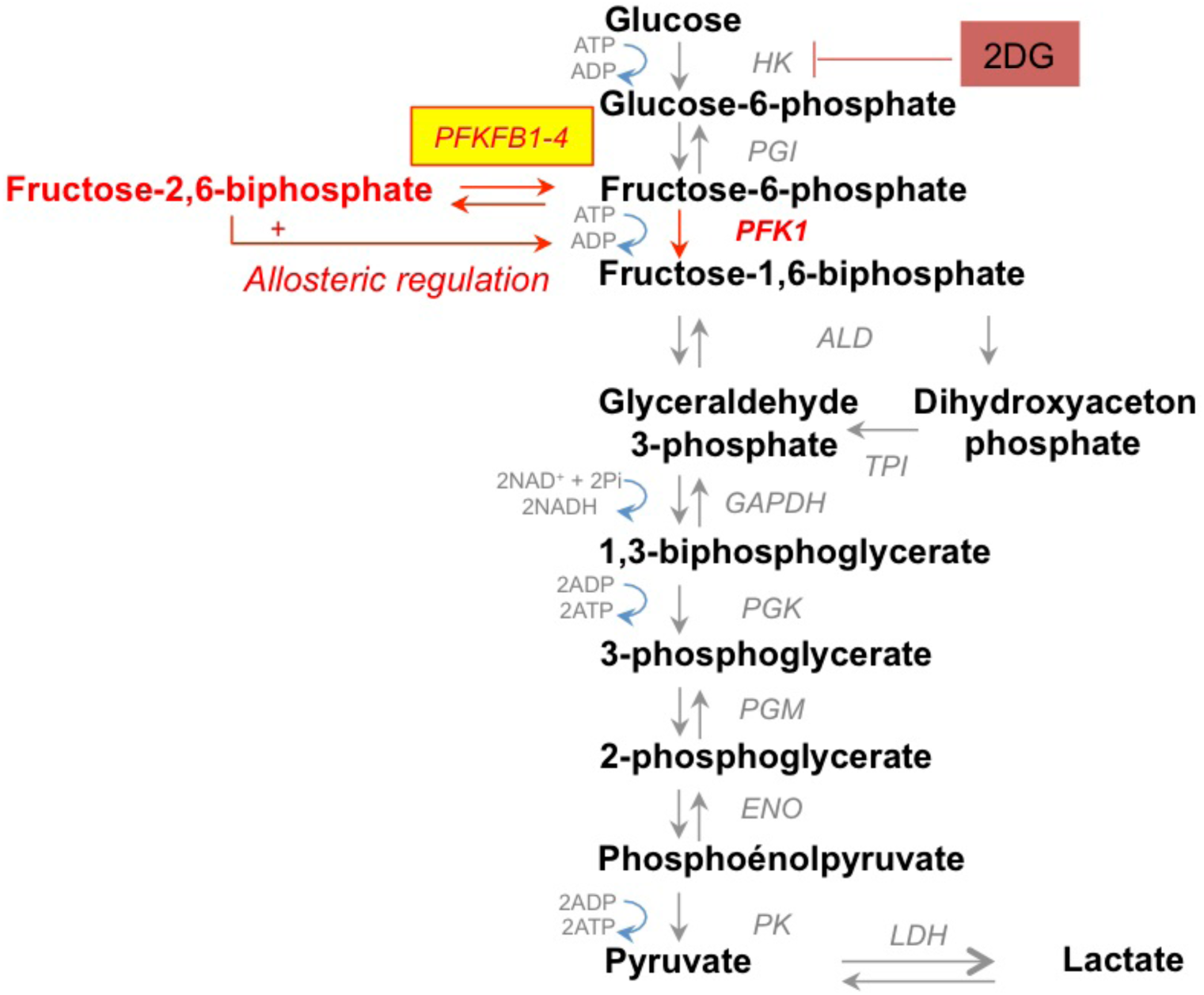
Schematics of glycolysis. Glucose phosphorylation by hexokinase is the first irreversible step of glycolysis. The second and rate-limiting step of glycolysis is catalyzed by PFK1. This step is tightly controlled in cells, by the levels of Fructose-2.6-bisphosphate, potent allosteric regulator of PFK1. Fructose-2.6-bisphosphate is synthesized by PFKFB1-4 enzymes, which thus control the rate of glycolysis in the cells. PFKFB3 and PFKFB4, with a strong kinase activity over their phosphatase activity, promote Fructose-2.6-bisphosphate synthesis and stimulate glycolysis. In addition to their classic role in glycolysis regulation, recent unconventional roles of the PFKFB enzymes have been described in cancer or development (see text).

**Fig. S2.**
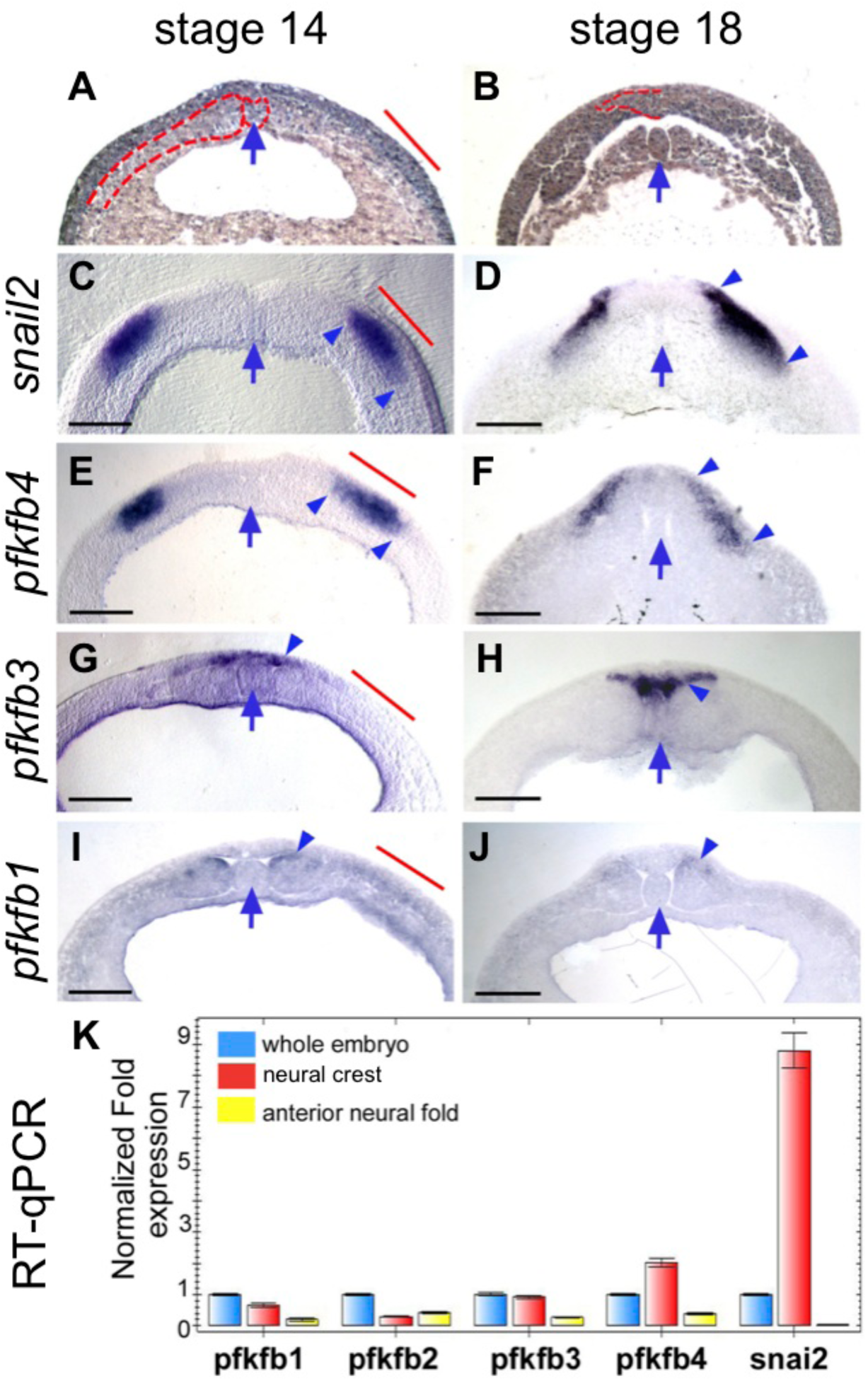
*Pfkfb4* is specifically upregulated in the neural border and neural crest during neurulation. (A-J). Transverse sections through the anterior neural plate at mid and late neurula stages (stage 14 and 18, respectively) allow comparing *Snail2* and *Pfkfb4* expression in the neural crest (*Snail2* and *Pfkfb4*), neural plate (*Pfkfb3*) and somites (*Pfkfb1*). (A,F): The notochord, neural plate and paraxial mesoderm are outlined on hematoxylin-eosin stained paraffin sections. (B-E,G-J) Vibratome sections of WISH-stained embryos. The arrow indicates the midline and the notochord. The blue arrowheads indicate in situ hybridization staining position. Red bar positions the neural border at stage 14. Scale bar = 200μm. (K) A quantitative analysis of *Xenopus laevis Pfkfb1*-*4* expression levels shows that *Pfkfb4* is specifically enriched at the neural border of frog neurulas (red), compared to the anterior neural fold (yellow). All values are normalized to *Odc* expression and to the average expression in a sibling whole embryo lysed at late neurula stage 18 (blue, value = 1). *Snail2* expression is used to monitor the quality of dissected tissues and shows high and specific expression at the neural border.

**Fig. S3.**
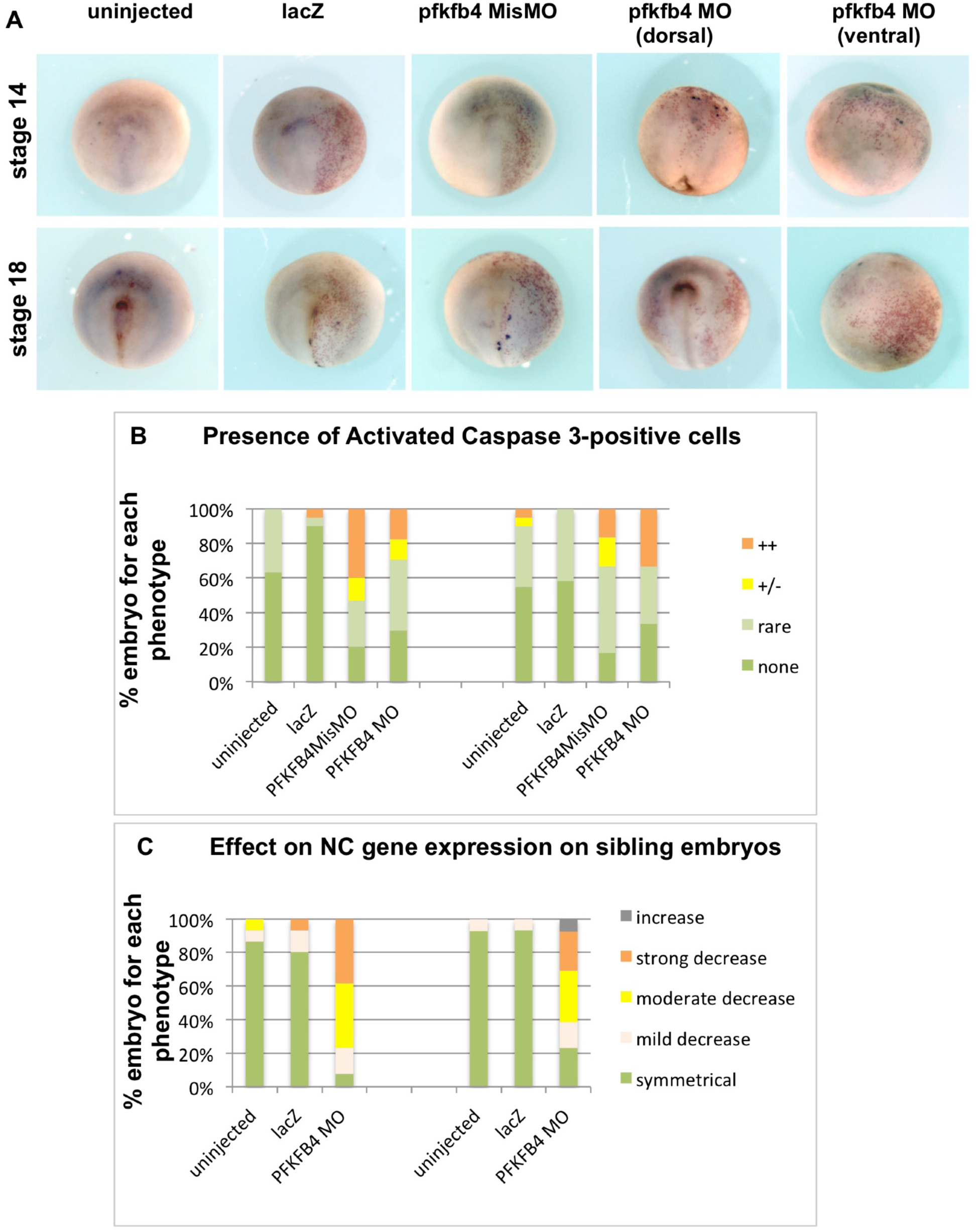
Patterning defects in NC are not due to increased cell death, after low-level PFKFB4 depletion in vivo. In this study, we use a low-level depletion of PFKFB4, with reduced MO concentrations, in order to bypass the earlier developmental arrest observed in previous study, when PFKFB4 severely depleted (above 60% mRNA depletion) and prevents gastrulating ectoderm specification (Pegoraro et al., 2015). Importantly, this severe depletion was rescued by adding back a MO-resistant *pfkfb4* mRNA, assessing both MO specificity and lack of non-specific toxicity (Pegoraro et al., 2015). (A) Here, we check that the morphant cells do not undergo elevated cell death at later stages of development by analyzing mid-neurulas at stage 14 and the end of neurulation stage 18. (B-C) We find less than 30% of embryos with caspase-positive cells in the lacZ-injected area (B), in conditions when over 70% of sibling embryos show deficient *twist* expression (C, either “strong”, “moderate” or “mild” decrease). The number of caspase-positive cells is slightly more elevated than in uninjected or lacZ-only-injected embryos, or in ventrally-injected morphants, but it remains limited to few cells and cannot account for the incidence of the patterning defects. Representative exemples of the majoritary phenotypes are shown for each condition, i.e. “rare” or “+/-” phenotypes. Embryos analyzed in this experiments: uninjected, n=49; lacZ-injected, n=46; PFFB4misMO-injected, n=16; PFKFB4MO-injected, n=43.

**Fig. S4.**
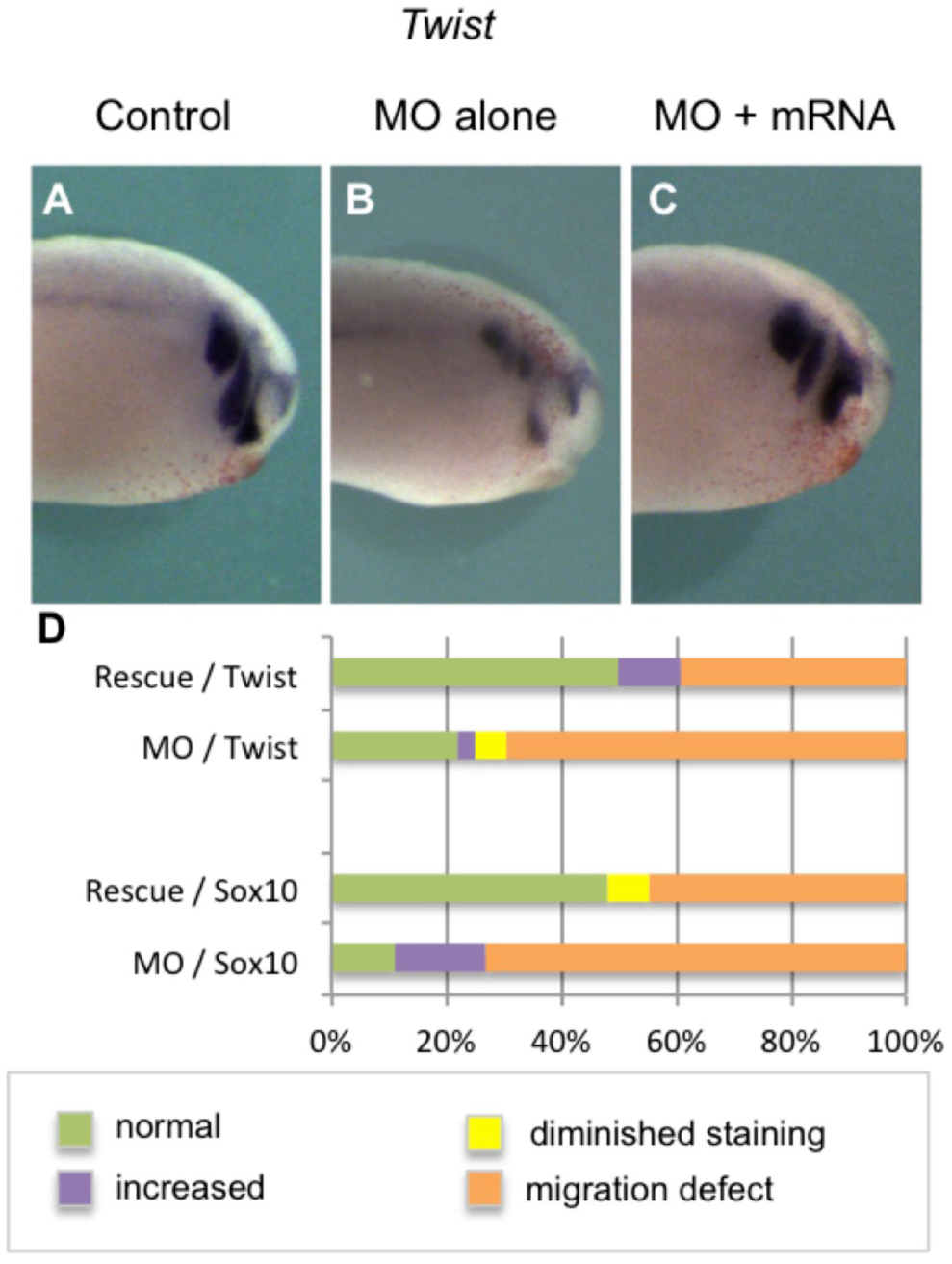
PFKFB4 is required for neural crest migration. (A,D) WISH analysis of neural crest migration at tadpole stages. When *Pfkfb4* MO is injected, embryos display decreased *Twist* expression and neural crest migration defects (B), when compared to control embryos (A). (C) In contrast, the co-injection of *Pfkfb4* mRNA with *Pfkfb4* MO restores *Twist* expression and NC migration, showing that the neural crest migration defects are due to PFKFB4 knowckdown. A-C are side views, anterior is to the right.

**Fig. S5.**
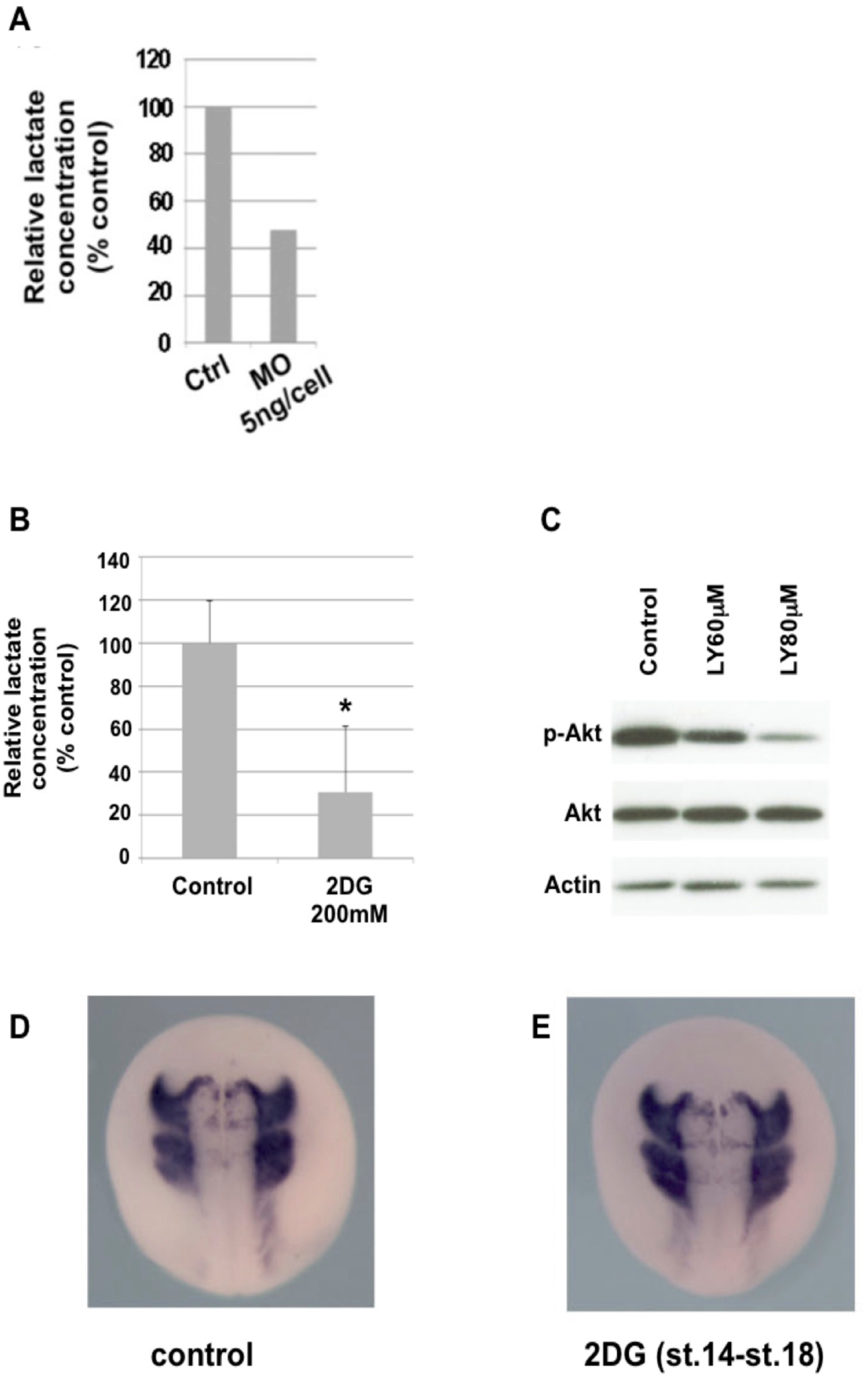
PFKFB4 depletion and 2DG treatment block glycolysis, LY294002 blocks AKT phosphorylation. (A) PFKFB4 low-level depletion affects lactate levels in embryos, indicating efficient glycolysis blockade. (B, C) Lactate concentration and Akt phosphorylation were decreased following 2-DG or LY294002 treatment (LY, two doses 60-80 μM), respectively. Error bars represent s.e.m (t-test p<0.05). (C,D) Blocking glycolysis during neurulation, between st. 14 and st. 18 does not affect *twist* expression *in vivo*. (D) control and (E) 2DG-treated embryos displayed normal *twist* expression (n=15 each).

**Fig. S6.**
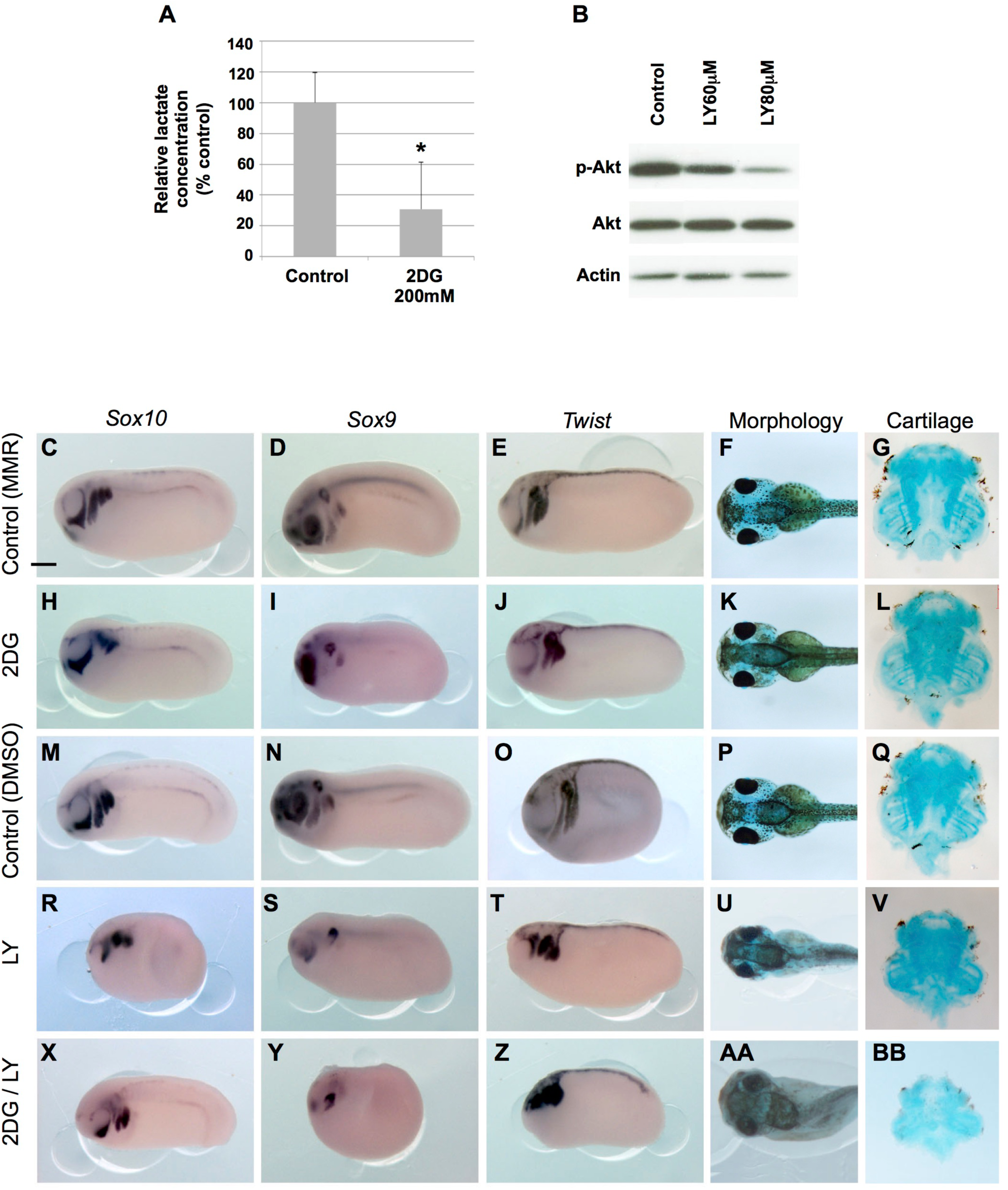
Glycolysis and PI3K-Akt signaling control neural crest migration. Glycolysis blockade (A-J) and PI3K-Akt blockade (K-T) during EMT and neural crest migration (stage 18 to stage 24) affects neural crest migration as shown by the expression of *Sox9*, *Sox10* and *Twist*. Embryos rinsed at stage 24 and grown until stage 45 display craniofacial and eye development defects, albeit the presence of all individual cartilage elements. (A-C,F-H,K-M,P-R, U-W) Side views, anterior is to the left. (D,I,N,S,X) dorsal views st 45. (E,J,O,T,Y) Ventral views of alcian blue stained visceral cartilages, st 45. Scale bar= 500μm.

**Fig. S7.**
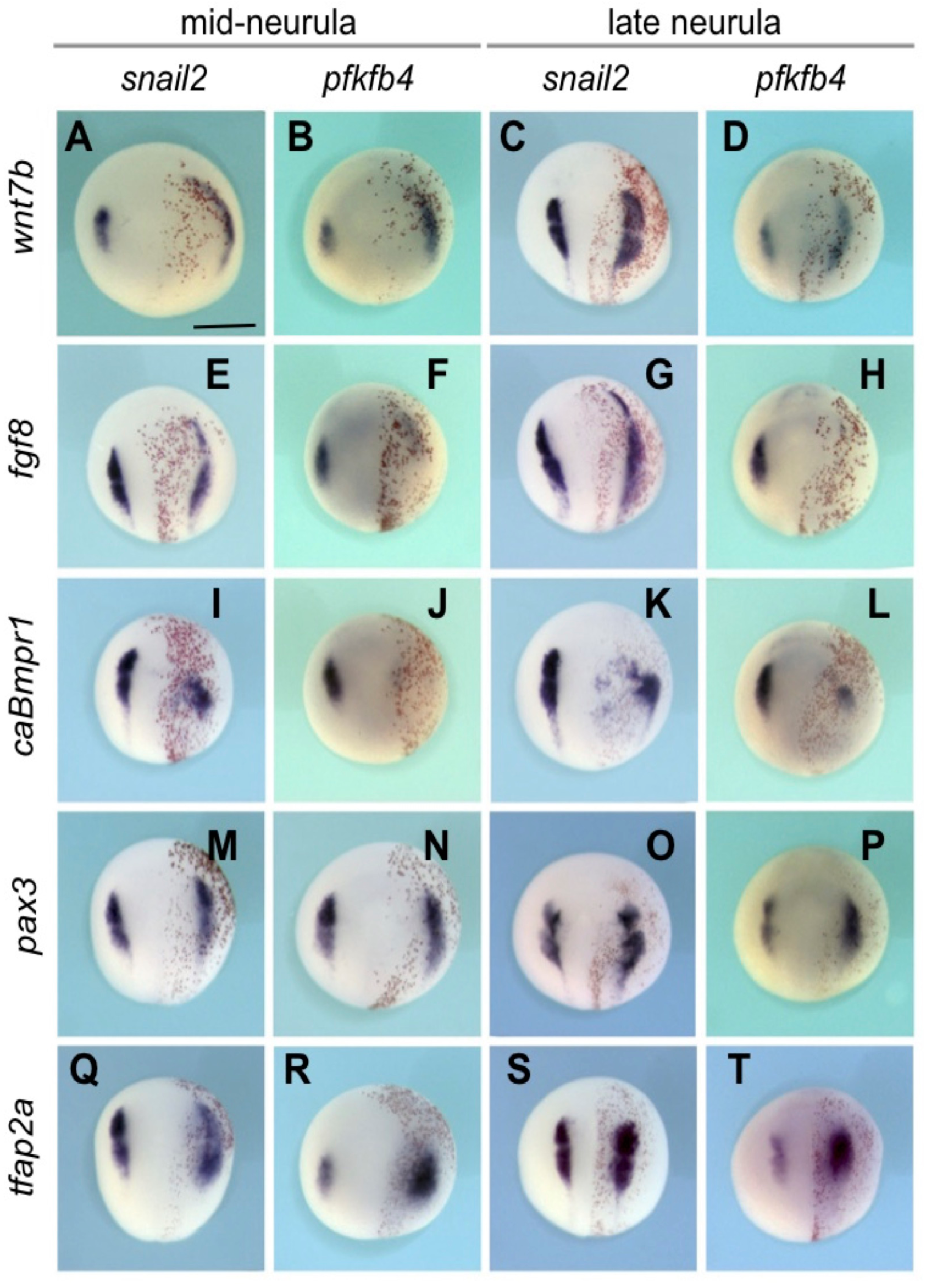
*Pfkfb4* is regulated by the neural crest gene regulatory network. Unilateral overexpression of members of the neural crest gene regulatory network modulates *Pfkfb4* expression at mid and late neurula stages (stage 14 and 18, respectively). (A-D) Overexpression of WNT signals? increases both Snail2 and Pfkfb4 expression. (E-H) FGF8 overexpression results in complete loss of *pfkfb4*, while *snai2* is expanded. (I-L) BMP overexpression with a constitutively active form of BMPR1 results in loss of *Snail2* and *Pfkfb4* expression in the NC and ectopic expression in the neural plate. (M-T) PAX3 and TFAP2a positively regulate *Pfkfb4* expression, similarly to *Snail2*. Dorsal views, anterior is to the top. Scale bar = 500μ.m.

